# A platform for efficient establishment, expansion and drug response profiling of high-grade serous ovarian cancer organoids

**DOI:** 10.1101/2022.04.21.489027

**Authors:** Wojciech Senkowski, Laura Gall-Mas, Matias Marin Falco, Yilin Li, Kari Lavikka, Mette C. Kriegbaum, Jaana Oikkonen, Daria Bulanova, Elin J. Pietras, Karolin Voßgröne, Yan-Jun Chen, Erdogan Pekcan Erkan, Mia Kristine Grønning Høg, Ida Marie Larsen, Tarja Lamminen, Katja Kaipio, Jutta Huvila, Anni Virtanen, Lars Engelholm, Pernille Christiansen, Eric Santoni-Rugiu, Kaisa Huhtinen, Olli Carpén, Johanna Hynninen, Sampsa Hautaniemi, Anna Vähärautio, Krister Wennerberg

## Abstract

The broad research use of organoids from high-grade serous ovarian carcinoma (HGSC) has been hampered by low culture success rates and limited availability of fresh tumor material. Here we describe a method for generation and long-term expansion of HGSC organoids with efficacy markedly improved over previous reports (55% vs. 23-38%). We established organoids from cryopreserved material, demonstrating the feasibility of using viably biobanked tissue for HGSC organoid derivation. Genomic, histologic and single-cell transcriptomic analyses revealed that organoids recapitulated genetic and phenotypic features of original tumors. Organoid drug responses correlated with clinical treatment outcomes, although in culture conditions-dependent manner and only in organoids maintained in human plasma-like medium (HPLM). Organoids from consenting patients are available to the research community through a public biobank and organoid genomic data explorable through an interactive online tool. Taken together, this resource facilitates the application of HGSC organoids in basic and translational ovarian cancer research.

## INTRODUCTION

High-grade serous ovarian carcinoma (HGSC) is the most prevalent and lethal type of ovarian cancer (OC), accounting for 70-80% of OC mortality^1^. HGSC is characterized by high molecular heterogeneity and shows only a few recurrent genetic abnormalities, including an almost universal loss of functional *TP53* (91-96% of patients) or mutations in *BRCA1/2* genes (20%)^2, 3^. HGSC patient survival rate has seen little improvement over the last few decades^1^. Cytoreductive surgery combined with platinum- and taxane-based chemotherapy remains the first-line treatment and despite the favorable initial response, most patients eventually relapse^1^. Introduction of poly(ADP-ribose) polymerase (PARP) inhibitors has increased the overall survival in patients with *BRCA1/2* mutations or other defects in DNA double-stranded break repair, highlighting that identification of predictive biomarkers for patient stratification can yield clinical benefits in HGSC^4^. Thus, capturing the enormous molecular complexity of HGSC in preclinical model systems has been deemed crucial to facilitate the discovery of new biomarkers and matched treatment strategies^1, 5, 6^.

Cancer organoids – patient-derived, self-renewing three–dimensional cell cultures – retain the genetic heterogeneity and recapitulate morphological characteristics of original tumors more closely than standard cell lines^7^, ^8^. They are also less costly, more scalable and easier to maintain than patient-derived xenografts (PDX). In recent years, organoids from OC and other solid tumors have been generated and utilized in molecular biology research and drug screening^9, 10, 11, 12, 13, 14^. However, organoid establishment success rates vary across tumor types, limiting their broad usability^15, 16^. Several studies in recent years have described short-term culture of primary HGSC cells (often, somewhat inaccurately, referred to as organoid cultures) with very high success rates^17, 18^. While establishment of organoids from many OC types has been successful, reported success rates of derivation of self-renewing, robust organoid cultures from HGSC have, in fact, remained relatively low, from 23% to 38%^11, 19, 20^. Furthermore, HGSC organoids have mainly been developed from fresh surgical specimens, which are viable only for a limited time and require geographical closeness and costly infrastructures between hospitals and research institutions. This poses major limitations for the establishment and broad availability of HGSC organoids for ovarian cancer research.

### Design

Cancer organoid culture media compositions vary across tumor types. Historically, most cancer organoid media were designed by altering the composition of pre-existing formulations used for maintenance of organoids from matching healthy tissue types. This method has been successful in the culture of cancer organoids from a number of tumor types, including low-grade serous, mucinous or mucinous borderline OC^11, 21^. However, in case of HGSC, this approach is sub-optimal, as cancer organoid media based on fallopian tube or endometrial organoid media were not sufficient to maintain the majority of attempted samples^11, 19^. Medium components promoting survival of healthy epithelium might also result in persisting contamination of organoid culture with normal cells, which has been observed in HGSC organoids^11, 19^. Furthermore, the physiological relevance of standard organoid media in functional assays has also been questioned. Organoid media are nutrient-rich and supplemented with a number of growth-promoting molecules, which could result in exaggerated growth rates and distorted drug responses^6, 22, 23^. Thus, there is a need to design medium conditions for efficient, long-term culture of HGSC organoids, but also to evaluate their relevance in organoid-based functional assays.

Here, we developed and optimized two novel medium formulations for long-term culture and expansion of HGSC organoids. With the new method, we generated a collection of 18 expandable HGSC organoid cultures from 11 patients, encompassing samples from different tissue sites and disease progression stages. We established all organoid cultures from cryopreserved samples, from 55% of attempted patients, a success rate markedly improved over previous reports. We validated the organoids using whole-genome sequencing (WGS), immunohistochemistry (IHC) and single-cell RNA sequencing (scRNA-seq) showing that they are genetically and phenotypically representative of the original patient samples over long-term culture. Based on patient consents, we deposited three organoid cultures in a publicly accessible biobank. We also observed that organoid drug responses in physiologic human plasma-like medium (HPLM) were markedly different from those in nutrient- and growth factor-rich expansion medium and were more closely correlated to clinical patient outcomes.

## RESULTS

### Establishment of a novel HGSC organoid culture medium

For HGSC organoid medium optimization and subsequent organoid derivation, we used cancer patient material from debulking surgery, laparoscopic biopsies or ascites drainage (Table S1). Following the surgical procedure, the tissue was immediately processed and frozen. Cryopreserved cell suspensions were shipped to the laboratory, thawed and seeded for organoid growth. Sample/organoid names indicate the patient number, clinical course phase at sampling and tumor location (for instance, EOC989_iOme = sample from patient EOC989, taken during interval debulking surgery, from tumor located in omentum; full explanation of abbreviations is available in Table S1).

As a starting point for medium optimization, we used Advanced DMEM/F12 medium, supplemented with Glutamax, Primocin, N-acetyl-cysteine and B27 Supplement (=‘Basal Medium’) (Figure 1A and 1E). We then assessed the influence of individual medium additives, reported by Hill *et al*., on the short-term organoid formation and growth of HGSC primary cells^17^. Addition of FGF-10, p38 inhibitor SB202190 or TGFβ receptor inhibitor A83-01 resulted in improved organoid formation (Figure S1). On the other hand, addition of EGF, FGF-2 or Noggin resulted in increased cellular attachment and decreased organoid formation. R-Spondin 1, nicotinamide (1 mM) or prostaglandin E did not cause any observable effect. Thus, we proceeded with the Basal Medium, supplemented with FGF-10, SB202190 and A83-10 (=’Medium 0.1; M0.1’) to further optimization (Figure 1A and 1E).

**Figure 1.**
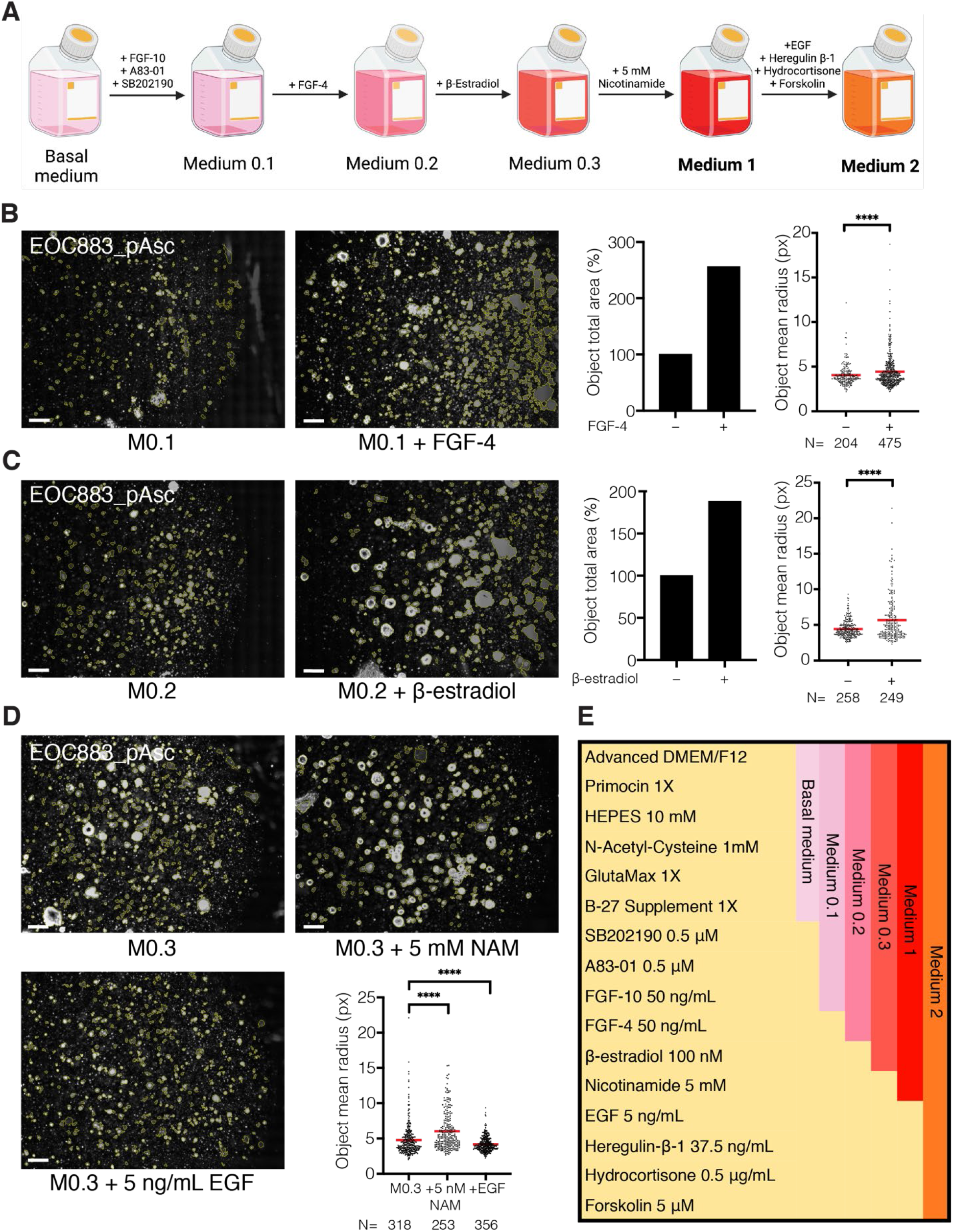
Establishment of new HGSC organoid media formulations. **(A)** Summary of the medium composition establishment process. **(B)** *Left:* Phase-contrast images of basement membrane extract (BME) droplets with objects (outlined in yellow) identified with CellProfiler. EOC883_pAsc cells were cultured in M0.1 or M0.1 supplemented with FGF-4 (10 ng/mL) for 38 days (passaged once on day 17). Scale bar, 200 μm. *Right:* Total area of objects and mean (marked with a line) object radius in the particular picture, estimated using CellProfiler. **(C)** *Left:* Phase-contrast images of BME droplets with objects identified as above. EOC883_pAsc cells were cultured in M0.2 or M0.2 supplemented with β-estradiol (100 nM) for 39 days (passaged once on day 20). Scale bar, 200 μm. *Right:* Total object area and mean object radius in the particular picture, as above. **(D)** *Top and bottom-left:* Phase-contrast images of BME droplets with objects identified as above. EOC883_pAsc cells were cultured in M0.3 or M0.3 supplemented with nicotinamide (NAM, 5 mM) or EGF (5 ng/mL) for 38 days (passaged once on day 19). Scale bar, 200 μm. *Bottom-right:* Mean object radius in the particular picture, as above. **(E)** Overview of particular HGSC organoid media formulations. **** = p<0.0001, unpaired two-tailed t-test.

Despite promoting growth after seeding, Medium 0.1 did not sustain growth over passaging. Thus, we set out to explore molecules that support the establishment of long-term, self-renewing organoid culture. We tested medium additives that have been reported to support cancer organoid growth as well as additives that have not been used as cancer organoid media components, but are important signaling molecules for cancer stem-like cells (CSCs) in HGSC. The full list of tested additives and their effects on HGSC organoid growth is available in Table S2. Of these, addition of FGF-4 resulted in increased organoid formation and growth over passaging (=’Medium 0.2; M0.2’; Figure 1B). Notably, FGF-4 has been previously reported to promote the tumorigenicity of OC CSCs^24^, but has not been included in cancer organoid media before. Further, we observed that addition of Wnt-pathway activating R-spondin-1 and Wnt-conditioned media, alone or in combination, caused decreased growth and HGSC organoid formation (Figure S2A). Wnt signaling inducers are essential components of media used for culturing organoids from normal epithelia^25, 26^, but multiple reports have indicated that they often are redundant in cancer organoid cultures^23, 27, 28^. Detrimental effects of Wnt on HGSC organoid culture have been previously described ^20^. However, other HGSC organoid media included Wnt-activating molecules in their composition^11, 17, 19, 21^.

In the following experiment, supplementation of M0.2 with β-estradiol increased the HGSC organoid formation and growth over passaging (=’Medium 0.3; M0.3’; Figure 1C), in concordance with previous reports^11,19^. Addition of nicotinamide further improved the organoid formation (Figure 1D), but the effect was concentration-dependent (Figure S2B). Addition of 5 mM nicotinamide was most advantageous, in agreement with the previous report by Maenhoudt et al.^19^, whereas 1 mM and 10 mM nicotinamide, reported by others^11, 17, 20^ yielded sub-optimal organoid growth. Thus, we included 5 mM nicotinamide in the final HGSC organoid medium formulation (=’Medium 1; M1’; Figure 1A and 1E).

### Two medium formulations improve the success rate of HGSC organoid culture

Interestingly, when M0.3 was supplemented with EGF, a component of all previously published HGSC organoid media, we observed markedly reduced growth in the EOC883_pAsc sample and eventual collapse of the culture (Figure 1D). In contrast, EGF promoted growth of several other samples (Figure S2C, Table S2). Combining EGF with additives reported previously by others^11, 19^, including heregulin-1β, hydrocortisone and adenylyl cyclase activator forskolin, further improved growth and expansion of these samples (Figure S2D, Table S2). On the contrary, EOC883_pAsc growth and organoid formation were harmed by these supplements (Figure S2E, Table S2). The full list of tested medium additives and their influence on particular samples is available in Table S2. These experiments demonstrated that the addition of EGF, heregulin-1β, hydrocortisone and forskolin can either promote or restrict organoid growth, depending on the sample. Thus, we concluded that, to maximize the likelihood of successful organoid growth, every HGSC sample should be cultured in parallel in two different media – M1 and Medium 2 (M2; M1 supplemented with EGF, heregulin-1β, hydrocortisone and forskolin).

We tested this strategy by culturing three HGSC samples in parallel in M1, M2 and previously published HGSC organoid media. As expected, EOC883_pAsc could only be successfully cultured in M1 (Figure 2A). In other formulations, the growth was limited and followed by the eventual collapse of the culture (Figure 2B). In contrast, EOC382_pOme could be cultured in M2 and the formulations from Maenhoudt *et al*. and Hoffmann *et al*.^19,20^ (Figure 2A). Notably, in the two latter formulations, which both contain higher concentrations of EGF than M2, the growth was accelerated when compared to M2 (Figure 2C), suggesting that EOC382_pOme benefits from growth-factor-rich conditions. Finally, EOC136_pAsc failed to stably grow over passaging in any of the tested medium formulations (Figure 2A), but short-term growth was observed to the largest extent in M2 (Figure 2D). Taken together, these results demonstrated that culturing HGSC primary cells in parallel in M1 and M2 enables successful derivation of organoids from samples, which would otherwise fail if cultured using previously published methods.

**Figure 2.**
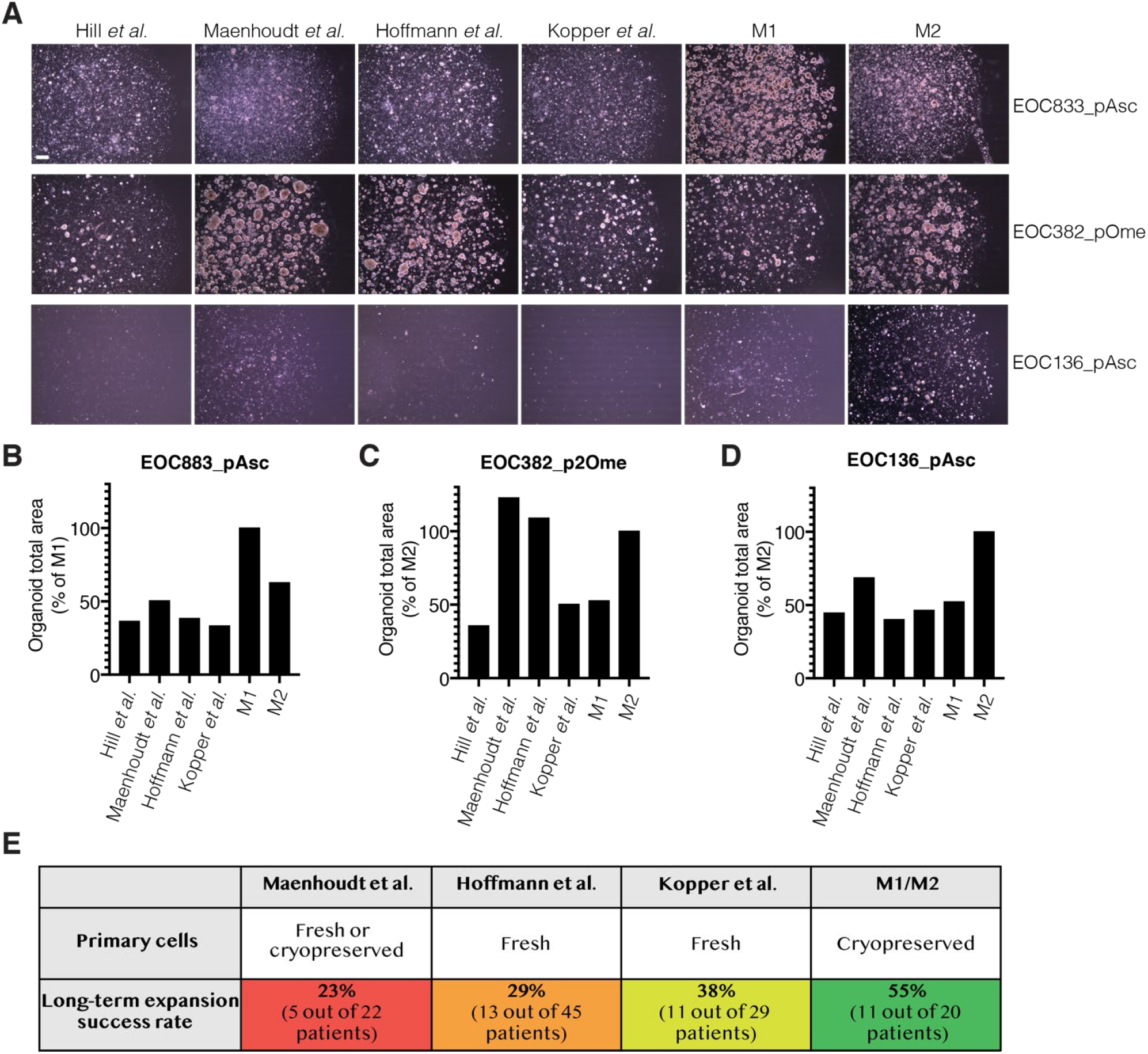
The M1/M2 method provides greater success rate of HGSC organoid culture than previously published protocols. **(A)** Phase-contrast images of EOC883_pAsc, EOC382_pOme and EOC136_pAsc cultures in previously published HGSC organoid media or M1/M2. EOC883_pAsc, EOC382_pOme or EOC136_pAsc cells were cultured for 36 days (passaged once on day 20), 31 days (passaged once on day 17) or 51 days (passaged once on day 29), respectively. Scale bar, 200 μm. **(B)** to **(D)** Total area of objects identified with CellProfiler in images presented in (A), for EOC883_pAsc (B), EOC382_pOme (C) and EOC136_pAsc (D). **(E)** Comparison of HGSC organoid culture establishment success rates reported previously (for source of the data in relevant publications, see Table S3) to the success rate of M1/M2 organoid culture.

This conclusion is corroborated by comparing previously reported HGSC organoid long-term culture success rates to the success rate of M1/M2 culture. Hill et al. only attempted short-term cultures^17^ and we therefore excluded this report from the comparison. Maenhoudt et al.^19^ succeeded in culturing organoids from five different HGSC patients (defined by the authors as >4 passages), out of attempted 22 patients, resulting in an overall success rate of 23% (for data source in ^19^, see Table S3; Figure 2E). Hoffmann et al.^20^ successfully grew organoids for 13 out of 45 patients (min. 6 passages in all reported cultures), with a success rate of 28% (Table S3, Figure 2E). Kopper et al.^11^ reported successful development of organoids that had not shown growth arrest and reached at least passage 8 for 11 patients out of 29 attempted (success rate of 38%; Figure 2E, Table S3). We used even more stringent criteria and defined a successful organoid culture if four conditions were fulfilled: 1) we managed to grow the cells for at least 10 passages; 2) we did not observe growth arrest in the sample; 3) we expanded the cancer cells in the sample; 4) the cultured cells carried the same *TP53* mutation as the original sample. Overall, we attempted to culture organoids from cancer material of 20 different patients. Using the M1/M2 method, we derived organoid cultures for 11 patients, and achieved a success rate of 55% (Figure 2E). Interestingly, if we had used only the EGF-containing M2, we would only have been able to derive cultures from 6 patients with a success rate of 30%, similar to other reports. Further, we established all cultures from cryopreserved material, demonstrating the feasibility of using viably biobanked tissue for HGSC organoid derivation with a high success rate. Maenhoudt et al. also reported long-term HGSC organoid cultures developed from frozen tissue, but with a low success rate (2 cultures developed out of 11 attempted).

### Establishment and characterization of expandable long-term HGSC organoid collection

In total, we developed 18 stable, long-term HGSC organoid cultures – 8 cultures in M1 and 10 in M2 (Figure 3A), The time required to reach a phase of sustained expansion varied across the organoid cultures, ranging from 26 days for EOC677_rAsc to 328 days for EOC41_pOme, and on average was 103 days (Table S1), Importantly, even though we observed cell growth and 3D structure formation in samples from 19 out of 20 patients (Figure S3A), initial material expansion did not always result in a stable organoid culture. For instance, EOC136_iOme ceased to expand beyond passage 8 and the culture collapsed (Figure 3A). However, in most cases, samples that reached a stable passaging rate (every 9-16 days, at a ratio ranging from 1:2 to 1:6) expanded throughout the tested period (up to passage 20). All organoid cultures were cryopreserved and resumed growth after resuscitation (100% success rate, n=18; Table S1).

**Figure 3.**
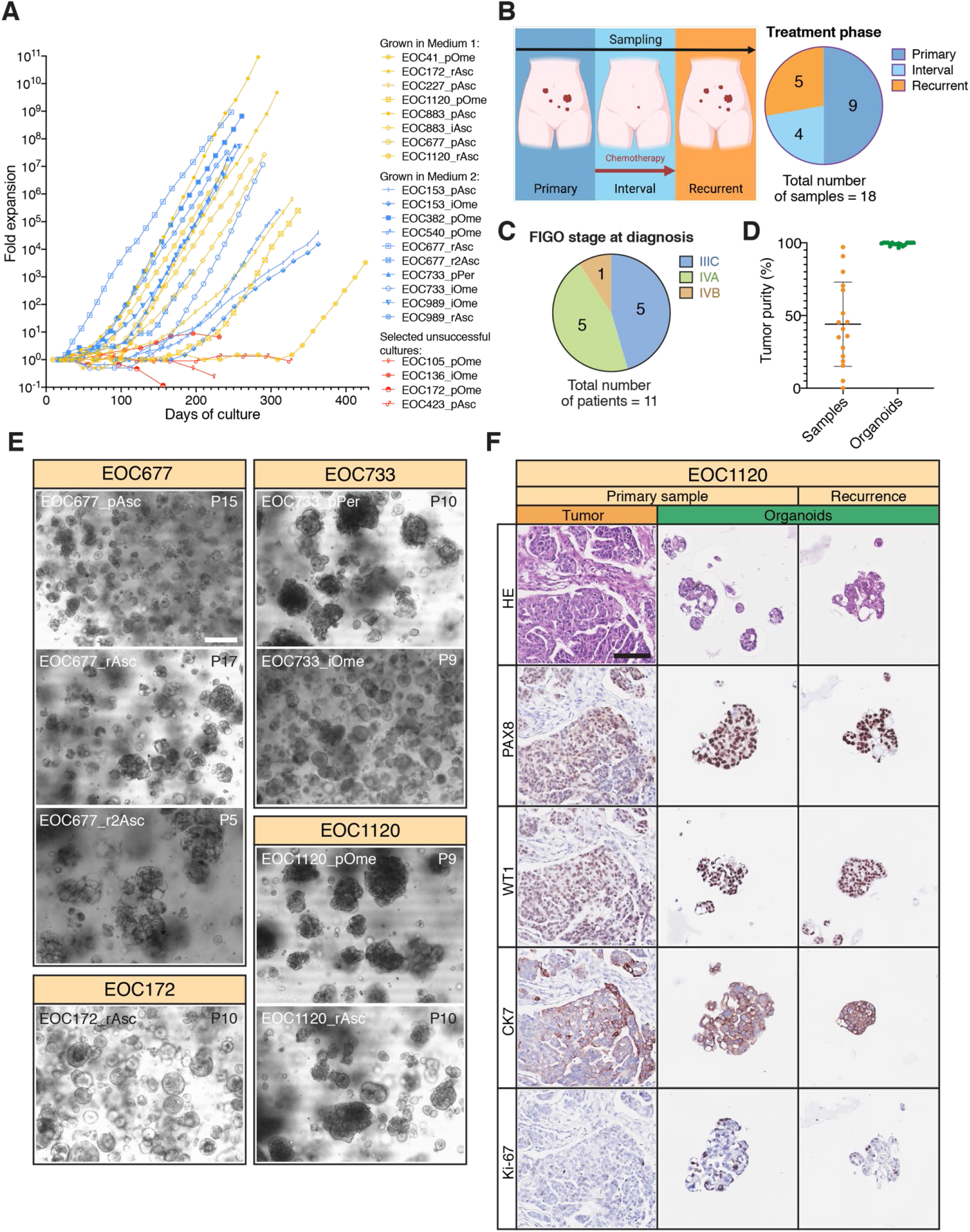
Overview of the HGSC organoid collection. **(A)** Growth curves of successful organoid cultures in M1 (n=8), M2 (n=10) and selected unsuccessful cultures (n=4). **(B)** and **(C)** Categorization of established HGSC organoid cultures according to the clinical course phase at sampling (B) and FIGO stage at diagnosis (C). **(D)** Tumor purity of tumor samples and corresponding organoid cultures presented as mean ± s.d. **(E)** Brightfield images of selected organoid cultures depicting various organoid morphologies. Passage numbers (P) indicated In top-right corners. Scale bar, 100 μm. **(F)** HE and IHC staining of EOC1120_pOme tumor tissue, organoids derived from it and organoids derived from relapsed tumor (EOC1120_rAsc). Organoids demonstrate morphological features similar to the original tissue, including nuclear pleomorphism, adenopapillary growth pattern and positive staining for PAX8, WT1 and CK7. They are also more proliferative than the original tissue, depicted by higher Ki-67 expression. Scale bar, 100 μm.

We derived organoids from patients sampled at different clinical course phases: before chemotherapeutic treatment (=primary, p’; n=9), during chemotherapy (=’interval, i’, n=4) and at relapse (=recurrence, r’, n=5) (Figure 3B, Table S1). For 6 patients, we were able to derive multiple organoid cultures. For instance, for patient EOC677 we developed a treatment-naïve organoid model (EOC677_pAsc) and organoid models from the first and second relapse (EOC677_rAsc and EOC677_r2Asc, respectively). Sampled patients were diagnosed at FIGO stages IIIC (n=5), IVA (n=5) or IVB (n=1) (Figure 3C, Table S1). Using WGS data, we estimated the cancer cell content of the organoids and the original samples. Organoids were characterized by high tumor purity (99.2±1.1%), contrary to the original samples they were derived from (44.1±29%) (Figure 3D). In contrast to earlier reports^11,19^, we did not observe any significant contamination with normal cells in any of the organoid models.

HGSC organoids exhibited broad morphologic heterogeneity in culture, at both inter- and intra-patient levels (Figure 3E). For instance, EOC677_pAsc organoids grew as small, densely-packed aggregates, while EOC677_rAsc and EOC677_r2Asc formed loosely-aggregated, cystic structures. Other observed structures include spheroid-like aggregates (for instance, EOC172_rAsc or EOC733_iOme) or large, irregular, densely-packed aggregates (for instance, EOC733_pPer, EOC1120_p¤me or EOC1120_rAsc).

To compare the organoids’ internal phenotypes to those of corresponding patient samples, we performed hematoxylin and eosin (H&E) and IHC stainings. Overall, organoids exhibited the morphological features of the matching tumor’s epithelium, including adenopapillary growth pattern and severe (3+) nuclear pleomorphism (Figure 3F, Figure S3A-S3E). All tested organoid-tissue pairs (n=5) were also concordant in the expression of HGSC IHC markers -paired box gene 8 (PAX8), Wilms’ tumor protein (WT1) and cytokeratin 7 (CK7) (Figure 3F, Figure S3A-S3E) – with organoids showing more homogeneous staining intensity and higher percentage of cells positive for each marker, compared with the more variable expression patterns in tumor tissue samples. Interestingly, IHC features of EOC1120_pOme tissue and organoids were preserved in organoids derived from the patient’s recurrent disease ascites (EOC1120_rAsc, Figure 3F, Figure S3E). As expected, organoid cells were relatively fast-growing and exhibited stronger staining (both in intensity and percentage of positive cells) for the cellular proliferation marker Ki-67 than cancer cells in the original tumor tissue (Figure 3F, Figure S3A-S3F).

### HGSC organoids retain the genomic landscape of patient samples over long-term culture

We performed WGS analysis to investigate whether the organoids recapitulated the genomic profiles of original patient tumors. All organoid models harbored *TP53* mutations matching those observed in corresponding tumor tissues (Figure 4A). Notably, the variant allele frequency (VAF) of *TP53* mutations was 1 for all organoid cultures, confirming that organoids comprised only cancer cells. Other genetic aberrations characteristic for HGSC included amplification of *CCNE1, KRAS, MYC* and *MECOM* and point mutations in *RB1, CSMD3, CDK12, KMT2B, KMT2C* and *CCNA2* (Figure 4A). In general, these were conserved between original tumor samples and organoids and over long-term passaging. The use of samples from different clinical course phases of the same patient enabled us to derive organoids representing tumors’ genetic evolution (for example, models with *de novo CDK12* mutation in patient EOC677, acquired at the first tumor recurrence) (Figure 4A). However, in some organoid cultures, we observed new mutations that appeared at late passages, suggesting genetic evolution or clonal selection during culture, as described in previous long-term organoid culture reports^9, 11^. Overall, organoids exhibited very high mutation concordance with the original patient samples, even when compared to control material from other tumor sites (Figure 4A and Figure S4).

**Figure 4.**
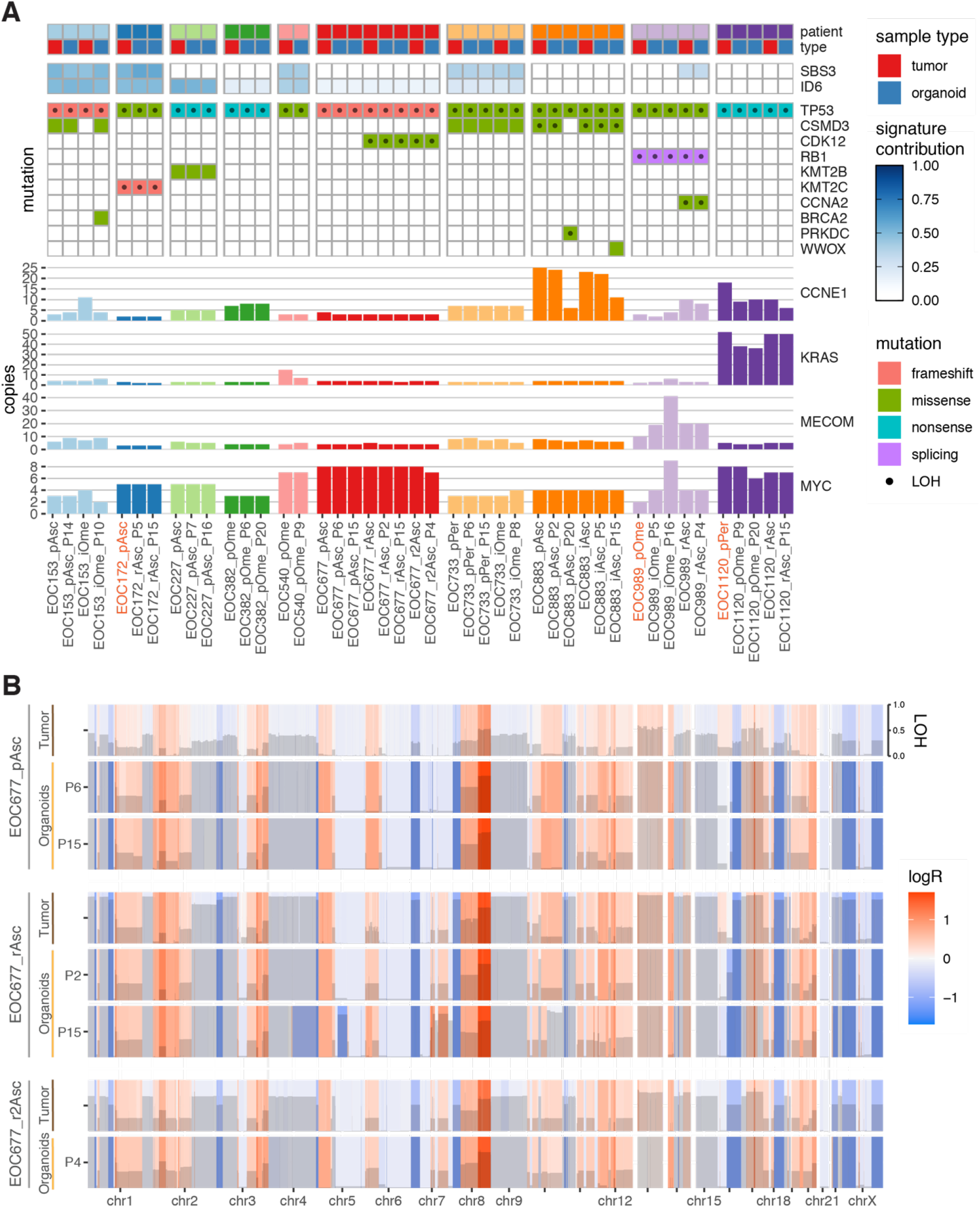
HGSC organoids cultured with M1/M2 recapitulate genomic landscapes of original tumor tissues over long-term culture. **(A)** Chart displaying somatic mutations and amplifications of selected, HGSC-relevant genes and contribution of HRD-associated mutational signatures (ID6, SBS3) in patient tumor tissue and corresponding organoid cultures. Passage numbers (P) at sequencing are indicated for organoid cultures. Sample names are typed in orange, where tumor tissue from a different metastatic location/clinical progression stage than the one used for organoid derivation is presented (due to limited matching tissue availability for sequencing). LOH – loss of heterozygosity. **(B)** Genome-wide CNV analysis of tumor tissue and corresponding organoids from patient EOC677, derived from material sampled at diagnosis, first and second recurrence. Copy number changes are expressed as logR and color-coded. The extent of LOH is displayed with gray bars. Passage numbers (P) at sequencing are indicated for organoid cultures.

Defects in the homologous recombination DNA-repair pathway are present in around 50% of HGSC cases. Alexandrov et al. have identified a number of mutational signatures associated with specific mutational processes (COSMIC v3.1), including single-base substitution signature 3 (SBS3) and small insertion and deletion signature 6 (ID6), that are associated with homologous recombination deficiency (HRD)^29, 30^. We fitted the COSMIC v3.1 signatures to mutational profiles of tumor samples and matching organoids and assessed their contribution to the overall mutational burden. SBS3 and/or ID6 signatures were identified in 12 out of 17 patient samples and their contributions were well reflected in matching organoid cultures (Figure 4A). Notably, organoids also recapitulated the emergence of SBS3 and ID6 signatures during clinical progression (for instance, SBS3 signature that emerged at recurrence in patient EOC989).

To compare genomic landscapes of organoids and matching tumor material, we performed copy number variation (CNV) analysis. Organoids maintained the CNV profiles of original tumors over long-term passaging (Figure 4B and Figure S5). In addition, organoids from different clinical course phases of the same patient maintained the genomic changes acquired during tumor evolution (for instance, gain in chromosome 7 fragment at the first recurrence in patient EOC677, Figure 4B). The only exception was the EOC153_iOme organoid culture, which was more similar to the primary tumor sample EOC153_pAsc than the interval deposit it was derived from (Figure S4). This culture also exhibited a point mutation in *BRCA2* gene (Figure 4A), detected neither in primary, nor interval tumor material, suggesting that organoids were derived from a small subpopulation of cancer cells. Overall, the present organoid collection recapitulated interpatient genomic heterogeneity and accurately mirrored disease evolution. To allow closer examination of the genomic landscapes, we provide the mutation, copy-number and signature profiles through an interactive visualization in the online tool GenomeSpy (https://genomespy.app/), accessible at: https://csbi.ltdk.helsinki.fi/p/senkowski_et_al_2022/.

### HGSC organoids recapitulate tumor’s single-cell transcriptomic heterogeneity

Transcriptomic profiling of HGSC organoids has only been performed on bulk cell mass before^11^. Thus, it has been unclear whether HGSC organoids represent patient tumors in terms of functional heterogeneity at the single-cell level. To address this, we performed scRNA-seq of 3 organoid samples – EOC883_pAsc, EOC382_pOme and EOC733_pPer – and the tumor/ascites material they were derived from. For patient EOC883, besides the ascites sample, we also profiled a primary adnexal tumor sample EOC883_pAdn. We performed unsupervised clustering on the total of 19,526 cells, resulting in 26 subclusters (Figure S5A). Cells from organoid samples formed organoid-line-specific clusters, while cells from patient samples formed multiple mixed clusters (Figure 5A). Marker expression analysis revealed that mixed clusters consisted of stromal or immune cells, while cancer cells formed patient-specific clusters (Figures 5B and 5C). As expected, all cells from organoids were categorized as cancer cells (Figure S5B). We then asked whether expression of genes identified as patient-specific markers in tumor samples was reflected in the corresponding organoids. We observed good concordance between the tumor samples and organoids of patients EOC883 (9/9 concordant markers) and EOC733 (7/9) (Figure 5D). In contrast, in EOC382, only 4 out of 15 patient-specific markers were concordant in the organoids. However, we found 7 out of the 11 non-concordant markers to be immediate-early genes (IEGs). IEG expression has been described to be rapidly induced by a number of stimuli, including protease treatment, tissue dissociation or inflammatory signals from the tumor microenvironment^31, 32, 33^. Of note, EOC382_pOme tumor sample preparation was extraordinarily harsh and involved multiple additional filtering and gradient purification steps, which could have induced additional cell stress. This suggests that comparison of expression profiles of tumor sample and organoids from this sample likely is influenced by the tissue processing or tumor microenvironment signaling. Overall, we observed a good correlation between the expression of patient-tissue-based and organoid-based markers for all 3 profiled patients (Pearson r = 0.671, 0.752 and 0.752 for EOC883, EOC382 and EOC733, respectively; p<10^-15^ for all) (Figures 5E, S5C and S5D).

**Figure 5.**
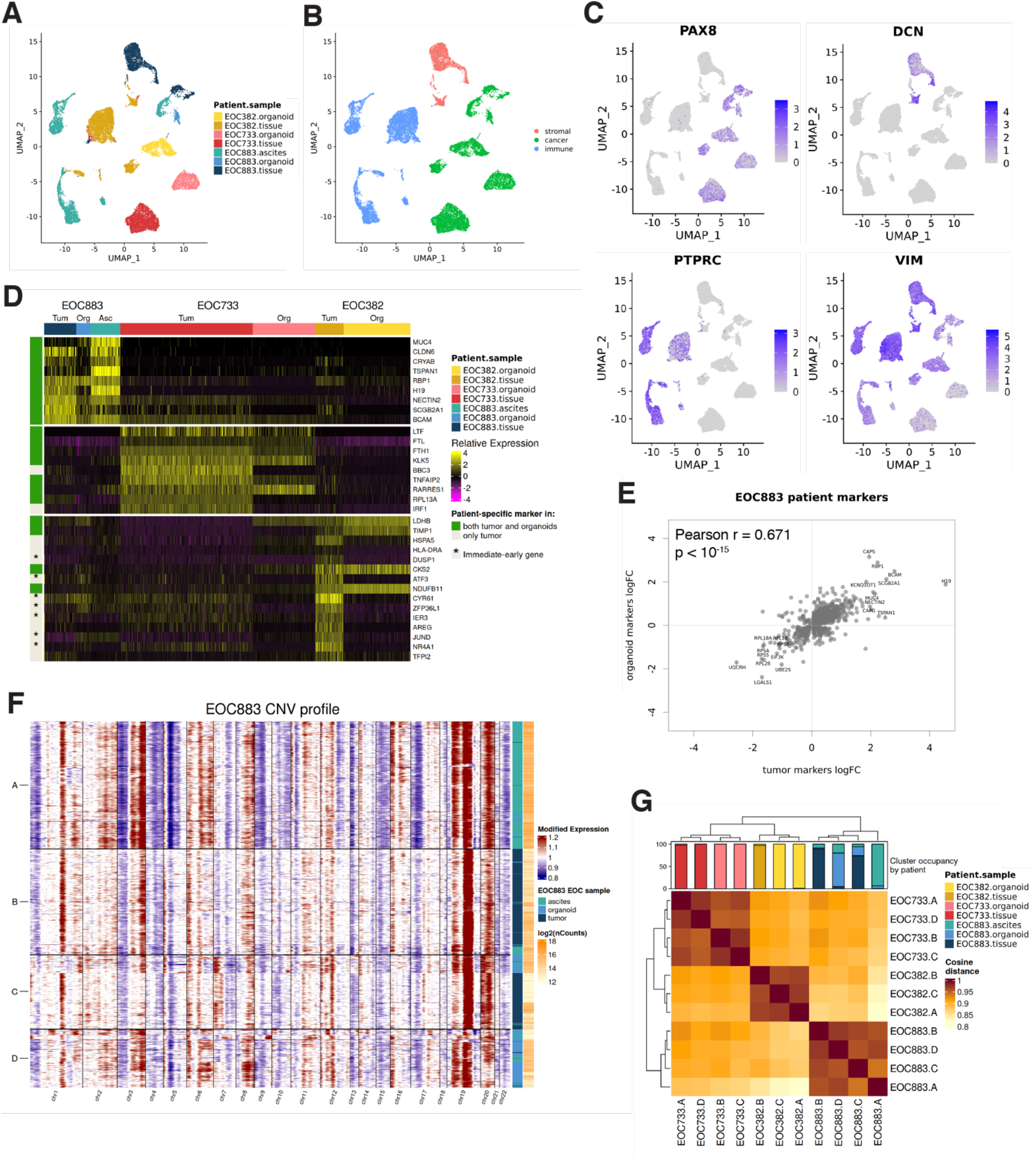
HGSC organoids preserve transcriptomic heterogeneity of original tumors. **(A)** UMAP visualization of 19,526 cells from tumor samples: EOC883_pAsc, EOC883_pAdn, EOC382_pOme and EOC733_pPer and corresponding organoids (except for EOC883_pAdn), with color-coded assignment to particular sample. **(B)** The same UMAP visualization with color-coded assignment to particular cell type (tumor, stromal or immune). **(C)** Single-cell expression of *PAX8, DCN, PTPRC* or *VIM* visualized in the same UMAP plot. **(D)** Heatmap displaying single-cell expression of genes that are most significantly overexpressed (“patient-specific markers”) in each clinical sample EOC883_pAsc, (Asc), EOC382_pOme (Tum) or EOC733_pPer (Tum) compared to single-cell expression of these genes in EOC883_pAdn (Tum) tumor sample and organoids (Org). Genes that are significant patient-specific markers in organoids in addition to the corresponding original tumor/ascites sample are marked with green rectangles. IEGs are marked with “*”. **(E)** Pearson correlation plot of patient-specific markers expression in EOC883_pAsc tumor sample and corresponding organoids. **(F)** Single-cell CNV plots from EOC883_pAsc tumor and organoids and from EOC883_pAdn tumor, inferred using InferCNV and classified into 4 subclusters. All analyzed samples are represented in each of the subclusters. **(G)** Heatmap displaying cosine distances between all subclusters in all samples, demonstrating patient-specific clustering of organoids and matching tissue samples.

Next, we investigated whether HGSC organoids represent the original patient samples at the cell subpopulation level. For this, we used the InferCNV^34^, which predicts genomic copy-number variation changes at a single-cell resolution and identifies subclusters of similarly altered cells within a population. Using InferCNV, for each patient we analyzed all tumor sample and organoid single-cell profiles as a single dataset and identified 3 to 4 main genetic subpopulations. For patient EOC883, all 4 subclusters contained profiles of all 3 analyzed samples (ascites, ascites-derived organoids and adnexal tumor tissue) (Figure 5F), indicating that organoids faithfully recapitulate the functional subpopulation composition of the original tumor. Subclusters identified for patients EOC382 and EOC733 were mixed to a lesser extent (Figures S6E and S6F). However, subpopulations from all patients, when analyzed as a single dataset, showed patient-specific clustering (Figure 5G). Taken together, these data indicate that organoids demonstrate a high degree of transcriptional similarity at the single-cell level to original tumor samples.

### HGSC organoid drug responses correlation to patient clinical outcomes is culture medium-dependent

Finally, we performed drug-response profiling of eight organoid cultures to investigate whether organoid drug responses correlate with those previously observed in patients. For this, we seeded organoid fragments, suspended in a basement membrane extract, into ultra-low attachment 384-well microplates and covered the cultures with sample-appropriate growth medium (Figure 6A). We and others have previously demonstrated that cell culture conditions, including non-physiologic concentrations of glucose, glutamine and other nutrients, can drastically influence drug responses in functional assays^35, 36, 37, 38^. Thus, to explore whether culturing conditions impact the correlation between organoid drug responses and clinical responses, we exchanged the growth medium to human plasma-like medium (HPLM) in half of the seeded plates after an initial period of organoid growth. HPLM mimics the metabolic composition of human plasma and has been shown in cancer cell lines to provide a more physiologically relevant environment for assessing drug responses^37^.

**Figure 6.**
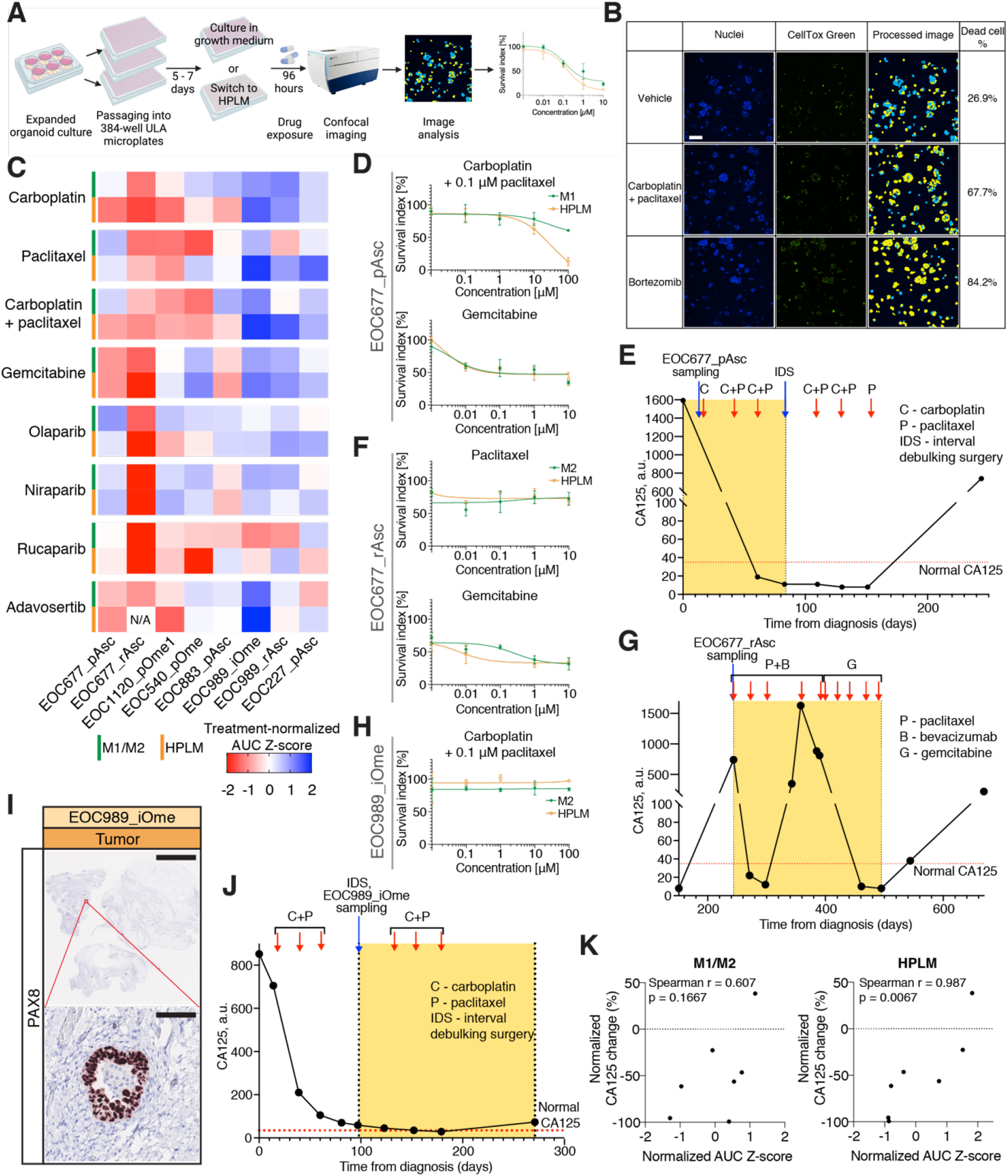
Correlation of HGSC organoid drug responses to clinical outcomes is culture medium-dependent. **(A)** Overview of the drug response profiling assay. **(B)** Image-based cytotoxicity assay. After drug exposure, cells’ nuclei in organoids are stained with Hoechst 33342 (blue, Nuclei) and dead cells are stained with CellTox Green (green). Organoids are imaged using automated confocal fluorescence microscopy, dead cell percentage is estimated using image analysis and normalized to negative (Vehicle) and positive (Bortezomib) control values. Scale bar, 100 μm. **(C)** Heatmap displaying treatment-normalized AUC Z-score values, showing responses of particular organoid cultures in growth medium (M1/M2) or HPLM to a panel of HGSC-relevant drugs. **(D)** Dose-response curves of EOC677_pAsc organoids treated with carboplatin + 0.1 μM paclitaxel or gemcitabine, in M1 or HPLM. Results are shown as mean ± s.d. (n = 2-3). **(E)** CA125 blood levels of patient EOC677 over time, during first-line therapy. Period relevant for comparison with *in vitro* drug response indicated with yellow rectangle. Normal CA125 range (<35 a.u.) indicated with a red dotted line. **(F)** Dose-response curves of EOC677_rAsc organoids treated with paclitaxel or gemcitabine, in M2 or HPLM. Results are shown as mean ± s.d. (n = 2-3). **(G)** CA125 blood levels of patient EOC677 over time, at first recurrence, second-line therapy. Data presented as in (E). **(H)** Dose-response curves of EOC989_iOme organoids treated with carboplatin + 0.1 μM paclitaxel, in M2 or HPLM. Results are shown as mean ± s.d. (n = 2-3). **(I)** IHC staining of EOC989_iOme tumor tissue for PAX8. Scale bars, 5 mm (*top*), 100 μm *(bottom)*. **(J)** CA125 blood levels of patient EOC989 over time, at first-line therapy. Data presented as in (E). **(K)** Spearman correlation plots between normalized AUC Z-score for carboplatin + 0.1 μM paclitaxel combination in organoids and normalized change (expressed as % of maximal patient-specific CA125 level in the relevant period) of blood CA125 in corresponding patients after carboplatin + paclitaxel chemotherapeutic treatment. N=7.

We exposed organoids in both growth and HPLM media to a panel of drugs used in HGSC clinical treatment (chemotherapeutics: carboplatin, paclitaxel, carboplatin/paclitaxel combination and gemcitabine; PARP inhibitors: olaparib, niraparib and rucaparib) and Wee1 inhibitor adavosertib, currently in clinical development for HGSC. We assessed the cytotoxicity by dead cell fluorescent staining and high-throughput confocal imaging (Figure 6B). Overall Z-factor across the experimental plates was 0.6, confirming the suitability of the assay (Figure S6A).

Organoids demonstrated differential responses to the drug panel (Figure 6C). Sensitivity of particular organoid cultures to gemcitabine, PARP inhibitors and adavosertib was similar in the M1/M2 and HPLM. However, responses to first-line chemotherapeutics carboplatin, paclitaxel and their combination often differed between media. For instance, organoids from the EOC677_pAsc sample were resistant to the combination of carboplatin and paclitaxel when exposed to the drugs in the growth medium, but sensitive in HPLM (Figure 6D). We then explored whether the *in vitro* drug responses matched those recorded in corresponding patients in the clinic. Clinically, patient EOC677 demonstrated response to carboplatin/paclitaxel combination, indicated by reduction of CA125 blood level (from 1593 to 11 a.u.) and its subsequent stabilization in the normal range (<35 a.u.) (Figure 6E). This corresponded to the organoid drug response in HPLM, but not in the growth medium. At relapse, the patient was treated with dose-dense paclitaxel, inducing a transient response followed by resistance to the therapy, which matched the EOC677_rAsc partial sensitivity to paclitaxel both in growth medium and in HPLM (Figures 6F and 6G). Subsequent treatment with gemcitabine induced longer-lasting normalization of CA125 and complete radiological response, which corresponded to sensitivity of organoids (notably, both EOC677_pAsc and EOC677_rAsc) in both media (Figures 6C, 6F and 6G).

Interestingly, we noted that EOC989_iOme organoids were fully resistant to the combination of carboplatin and paclitaxel (Figure 6H), while in the clinic, the patient EOC989 experienced CA125 level reduction after 3 courses of neoadjuvant carboplatin and paclitaxel. However, the sample, from which organoids were derived (Figure 6I), was acquired during the IDS after 3 cycles of chemotherapy, when CA125 was nearing the normal blood level (Figure J). Correspondingly, IHC staining of the EOC989_iOme tissue for PAX-8 revealed only small deposits of surviving cancer cells within the tumor (Figure 6I). Thus, as these cells gave rise to the EOC989_iOme organoid line, it is not surprising that we observed drug resistance *in vitro*. Accordingly, EOC989_rAsc organoids, derived from a relapsed sample from the same patient, exhibited resistance to carboplatin/paclitaxel combination (Figures S7B and S7C). These findings demonstrate that the disease clinical course phase at sampling and previous clinical history influence the comparison of *in vitro* to *in vivo* drug responses. Taking this into account, we asked if there was a correlation between sensitivity to carboplatin/paclitaxel combination and clinical outcome of chemotherapy (expressed as a percentage of CA125 blood level change in the period relevant for a particular sample/therapy). For the analysis, we chose only organoid cultures derived from samples acquired before the carboplatin/paclitaxel combination treatment (n=7, Figures 6D, 6E,6H, 6J, S6B-S6K). For organoids exposed to the combination in M1/M2, the correlation was moderate and not statistically significant (Spearman r = 0.607, p = 0.1667) (Figure 6K). In contrast, organoid drug responses in HPLM strongly correlated to CA125 reduction in corresponding patients (Spearman r = 0.987, p = 0.0067). Taken together, these results underscore the importance of medium conditions for the clinical relevance of organoid-based functional assays.

## DISCUSSION

Efficient establishment of long-term HGSC organoid cultures is essential for their broad application in OC research. Some studies have reported high success rates of short-term primary HGSC cell culture^17, 18^, but reported success rates of stably expanding HGSC organoid culture establishment were markedly lower^11, 19, 20^. In this context, our study confirms that a wide variety of culturing conditions allow for short-term HSGC cell survival and limited expansion from the majority of HGSC samples (in our case, samples from 95% patients attempted), but also that the long-term, robust HGSC organoid expansion is only possible under a restricted set of growth conditions. We present a novel method for HGSC culture, which enabled the generation of expandable HGSC organoid collection with a 55% success rate, markedly higher than in previous reports. We established all cultures from cryopreserved material, which is a major advantage over published protocols, as the common problem of the limited availability of fresh cancer tissue can be circumvented. Extensive testing of organoid culture media components resulted in two different medium formulations – M1 and M2. Eight samples in our collection grew in M1, which is relatively scarce in growth factors, compared with other previously published OC organoid media. Notably, unlike any previously published medium, M1 does not contain EGF, which we found to be harmful for some HGSC samples. In contrast, 10 samples were sustained by the growth-factor-rich M2. Furthermore, we found FGF-4, previously not used in cancer organoid culture, to be beneficial for HGSC organoid growth. These findings suggest that HGSC samples exhibit differential needs for sustained growth and that organoid media design should address this heterogeneity.

Importantly, HGSC organoids in our collection were pure cancer cell cultures, with >99% tumor purity based on CNV analysis and harbored the correct *TP53* mutation with VAF=1. We have not encountered any contamination with normal cells, a problem described in previous HGSC organoid reports^11, 19^. Presumably, this can be attributed to the lack of Wnt pathway stimulants in M1 and M2. Similarly, Hoffmann et al. found that HGSC organoids require low-Wnt environment, but organoid tumor purity is not defined in that report^20^. In contrast, Kopper *et al*. used Wnt-stimulating Rspo1 and, for many cultures, supplemented the medium with Wnt-conditioned medium. Tumor purity in HGSC organoids in that report ranged from 46% to 100% and, on average, was 90%^11^. Maenhoudt *et al*. also used Rspo1 and reported tumor purities in three organoid models ranging from 90% to 98%, and developed one organoid culture that eventually turned out to be derived from healthy tissue^19^. Together, these findings indicate that HGSC organoids require different niche factors than the matching healthy tissue and that supplementing the HGSC media with Wnt activators may provide a selective growth advantage to normal cells, resulting in decreased organoid tumor purity.

We established an organoid collection from samples acquired at different times of clinical progression. This enabled us to derive sequential organoid models from eight patients reflecting their clinical history–‘primary’ organoids representing treatment-naïve tumor; ‘interval’, from tumor cells that survived neo-adjuvant chemotherapy and organoids from the relapsed disease (‘recurrence’). This approach provides a unique collection of models that could be used to study differences in drug responses, changes in genetic dependencies and drug resistance mechanisms over disease progression, which are the most significant challenges in the HGSC patient treatment. Based on available consents from patients, we have deposited 3 organoid cultures – EOC733_pPer, EOC733_iOme and EOC41_pOme in a public biobank (Auria Biobank, Turku, Finland, https://www.auria.fi/biopankki/en/), from which they are available to the research community. Genomic data, including mutational, CNV and HRD-related signature profiles from all organoid cultures are available to explore through the online visualization tool GenomeSpy (https://csbi.ltdk.helsinki.fi/p/senkowski_et_al_2022/).

We characterized the organoid collection using WGS and HGSC-relevant IHC. Overall, organoids were representative of their tissue of origin and genetically stable over long-term passaging. The only exception was the EOC153_iOme line, which was more similar to the primary tumor than the interval sample. Genetic analysis demonstrated that it was likely derived from a small cell subpopulation that survived the selection pressure of the *in vitro* culture. This observation highlights the need for in-depth validation of all experimental models in organoid research. We also characterized three organoid cultures and their tissues of origin using scRNA-seq. The analysis demonstrated that HGSC organoids represent the phenotypic heterogeneity and display patient-specific expression features of the original tumors, indicating the suitability of the model for organoid-based functional assays.

We also observed major differences in organoid drug responses in different culturing media. When organoids were exposed to the drugs in the growth medium, drug responses only weakly correlated with clinical responses to the corresponding chemotherapeutics. This contrasts with previous reports, where strong correlations were found in HGSC^39^ and in other tumors^13, 40, 41, 42^. On the contrary, *in vitro* drug responses in HPLM correlated very well with *in vivo* drug responses, highlighting the impact of culture conditions on organoid drug responses. As cancer organoids have been proposed as predictive models for personalized medicine, our results suggest a need for large-scale evaluation of the physiological relevance of organoid cell culture conditions.

In summary, we present a new method for efficient HGSC organoid derivation and long-term culture. We provide a comprehensive validation of the models, demonstrating that they preserve the genetic and phenotypic heterogeneity of the original tumors. We also highlight the need for further evaluation of culture conditions to facilitate broad application of HGSC organoids in cancer research and their relevance in personalized cancer medicine.

### Limitations

Despite the success rate improved over previous methods, we were not able to establish long-term organoid cultures for 45% of patients. One reason for this may be poor viability of some cryopreserved samples upon resuscitation. However, it is likely that the pro-growth signaling needs of these samples are yet to be discovered and additional medium formulations could further increase the success rate of HGSC organoid culture establishment. Interestingly, we noted that successfully grown samples carry *CCNE1* amplification, a negative prognostic factor in HGSC^43^, more often than the failed-to-grow samples, indicating that this genetic aberration may facilitate cell growth *in vitro*. We were unable to find any other significant difference between the two groups, presumably due to a moderate number of established cultures.

## METHODS

### HGSC tumor samples and clinical data

The study used tumor material from patients who participated in the European Union’s Horizon 2020-funded HERCULES study (ID: 667403). All patients consented to collection, storage and research use of their tumor material and related data. The collection and storage of the material has been approved by The Ethics Committee of the Hospital District of Southwest Finland (ETMK): ETMK 53/180/2009 § 238. The patients were treated at The Department of Gynecology and Obstetrics at Turku University Hospital, Finland between 2016 and 2019. The patient outcomes were followed prospectively and their clinical outcome parameters such as platinum free interval (PFI) and response to chemotherapy were defined with RECIST 1.1 and GCIG criteria. Fresh tumor samples were obtained during tumor debulking surgeries, laparoscopic biopsies or ascites paracentesis. After acquisition, whenever possible, samples were divided in a few parts. A fraction of the sample was frozen at −150 °C for subsequent WGS and the rest was immediately processed.

### HGSC tissue processing

Solid fresh tissue samples were minced with scalpel and enzymatically digested overnight using 0.1X Collagenase/Hyaluronidase (#07912, Stemcell Technologies) in DMEM/F12 (#BE12-719F, Lonza) supplemented with 1X B-27 Supplement (#17504044, Gibco), 20 ng/mL hEGF (#E9644, Sigma) and 10 ng/mL FGF-b (#PHG0023, Gibco). Detached cells were purified with 70 μm and 40 μm filtering followed by gradient centrifugation with Histopaque-1077 (#10771, Sigma-Aldrich), according to the manufacturer’s instructions. Gradient centrifugation was repeated when necessary. Finally, the cells were washed with PBS, frozen in STEM-CELLBANKER (#11890, Amsbio) and stored at −150°C. Cell integrity and purity were assessed on cell smears on X-tra Adhesive Slides (#3800200, Leica) stained with toluidine blue (#89640, Sigma). Fresh ascites samples were centrifuged at 475 rcf for 15 minutes, followed by gradient centrifugation with Histopaque-1077 and freezing similarly to the solid tissue derived cells.

### Organoid media establishment experiments

Cryopreserved vials containing processed HGSC tumor material were shipped to the laboratory in dry ice and stored in liquid nitrogen until resuscitation for culturing. On the day of seeding, vials were taken out of the liquid nitrogen and thawed immediately in a 37°C water bath. Then, cell suspensions were mixed with minimal growth medium (depending on the medium establishment progress – Basal Medium, M0.1, M0.2, M0.3 or M1), that is: Basal Medium: Advanced DMEM/F12 (#12634010, Gibco), supplemented with 100 μg/mL Primocin (#ant-pm-1, Invivogen), 10 mM HEPES (#15630080, Gibco), 1 mM N-acetyl-cysteine (#A7250, Sigma), 1X GlutaMAX (#35050061) and 1X B-27 Supplement; M0.1: Basal Medium, supplemented with 0.5 μM SB202190 (#HY-10295, MedChemExpress), 0.5 μM A83-01 (#SML0788, Sigma) and 10 ng/mL recombinant human FGF-10 (#100-26, Peprotech); M0.2: M0.1, supplemented with 10 ng/mL recombinant human FGF-4 (#100-31, Peprotech); M0.3: M0.2, supplemented with 100 nM β-estradiol (#E2758, Sigma); M1: M0.3, supplemented with 5 mM nicotinamide (#N0636, Sigma. After mixing, cells were centrifuged at 200 rcf and washed with the minimal medium again. Next, cells were counted and viability was assessed using Countess Automated Cell Counter (Thermo Fisher). Then, cells were resuspended on ice in cold 7.5 ng/mL (concentration adjusted with cold PBS) Cultrex Reduced Growth Factor Basement Membrane Extract, Type 2, Pathclear (BME-2 #3533-010-02, R&D Systems) at a minimum of 10^6^ live cells/mL and seeded into pre-warmed 6-well cell culture plates (#140675, Nunc) in 20 μL droplets (10 droplets/well). Gel droplets were solidified in a humidified cell culture incubator at 37°C for 30-45 minutes and gently covered with 3 mL of the minimal or experimental culture medium (full list of tested supplements and manufacturers is available in Table S2). For the first 2-3 days after cell seeding, 5 μM Y-27632 (ROCK-1 and -II inhibitor, #HY-10583, MedChemExpress) was added to each of the media. Cultures were maintained in a humidified cell culture incubator at 37°C and culturing media were exchanged every 2-3 days. Cultures were regularly assessed using phase-contrast microscopy to monitor growth and organoid formation. Once pronounced growth was observed, cultures were photographed using a CCD camera (EC3, Leica) coupled to a phase-contrast microscope and passaged. Briefly, cultures were washed with PBS, covered with 2 mL of TrypLE Express (#12604013, Gibco), scraped off the cell culture plate surface and dissociated by vigorous pipetting. Then, the suspensions were incubated at 37°C for 15 minutes, transferred to 15 mL conical tubes and centrifuged at 300 rcf for 5 minutes. The supernatant was removed, cell pellets were resuspended in BME-2 and suspensions were seeded into 6-well culture plates and cultured as above. The passaging ratio for medium optimization experiments was 1:2. After 10-30 days of growth, cultures were photographed for image analysis.

### Image-based organoid growth assessment

All phase-contrast images used for the analysis were taken at 2.5X magnification. Photographs were analyzed using CellProfiler v3.1.9 ^44^. First, to make separate images comparable, illumination was equalized using “CorrectIllumination” modules with Gaussian filter. Subsequently, objects (organoids/cell clusters) were identified using “IdentifyPrimaryObjects” module in each image, within borders of the gel droplet (using a pre-prepared mask image for each analysis to define the droplet area). Objects were identified and single cells/large artifacts excluded from the analysis using Otsu three-class adaptive thresholding (typical diameter 20-400, adaptive window 150, smoothing 2, correction 1, smoothing filter 30). Then, total object area was estimated as a fraction of total area of all the objects divided by the area of the gel droplet. Object mean radius was also estimated for each object.

### Comparison of M1/M2 with previously published media

Previously published media were reproduced according to authors’ instructions. From Kopper *et al*. report, only the formulation without the Wnt-conditioned medium was used, due to previously observed negative impact of Wnt conditioned medium on HGSC samples. Cells from EOC883_pAsc, EOC382_pOme and EOC136_pAsc patient samples were resuscitated from frozen aliquots and seeded into 6-well culture plates and cultured as described above. 1 well/sample/medium was seeded, each containing 10 separate BME droplets. For the first 2-3 days after cell seeding, 5 μM Y-27632 was added to each of the media. Culturing media were exchanged every 2-3 days. EOC883_pAsc, EOC382_pOme or EOC136_pAsc cells were cultured for 36 days (passaged once on day 20, 1:1 ratio), 31 days (passaged once on day 17, 1:1 ratio) or 51 days (passaged once on day 29, 1:1 ratio), respectively. Then, cultures were photographed using a CCD camera (EC3, Leica) coupled to a phase-contrast microscope at 2.5X magnification and photographs analyzed as above. Organoid long-term culture success rate for HGSC in previous reports was calculated using published data (Table S3). Overall, all previously reported cultures were deemed long-term and expandable, unless stated otherwise. For the exact definition of a long-term, stable culture in each report, see the Results section.

### Organoid derivation and long-term expansion

Cells from patient samples were resuscitated from frozen aliquots and seeded into 6-well culture plates and cultured as described above. Number of wells and gel droplets seeded were dependent on the total number of live cells after resuscitation (10-60 droplets per sample). Each sample was initially cultured in M1 and M2 in parallel. Cultures were regularly assessed using phase-contrast microscopy to monitor growth and organoid formation. Once pronounced growth was observed, cultures were photographed using a CCD camera (EC3, Leica) coupled to a phase-contrast microscope and passaged. Initial passaging ratio was decided on a case-by-case basis, depending on the number of growing organoids observed. When very few organoids were observed, cultures were often initially passaged at 1:0.5-1 ratio, in order for organoids to reach higher density, which frequently resulted in increased expansion in subsequent passages. For densely growing cultures, passaging ratios were steadily increased until the cultures reached a stable expansion rate. In most cases, after 3-7 passages culture was continued only in M1 or M2, based on the observed growth and successful organoid formation. Cultures where no further growth was observed, were terminated. Overall fold-expansion was calculated as the initial number of seeded wells (counting 1 BME droplet as 0.1 well) multiplied by all previous passaging ratios. Organoids that reached a stable expansion rate were cryopreserved. Briefly, cultures were washed with PBS, covered with 2 mL of TrypLE Express (#12604013, Gibco), scraped off the cell culture plate surface and dissociated by vigorous pipetting. Then, the suspensions were incubated at 37°C for 15 minutes, transferred to 15 mL conical tubes and centrifuged at 300 rcf for 5 minutes. The supernatant was removed, cell pellets were resuspended in 1 mL of STEM-CELLBANKER and immediately frozen at −80°C. After 24 hours, the frozen vials were transferred to liquid nitrogen tanks for long-term storage. Organoid cultures were expanded until passage 20. Based on available biobanking consents, three organoid cultures – EOC4l_pOme, EOC733_pPer and EOC733_iOme – were deposited in Auria Biobank (Turku, Finland) and are available to the research community.

### Histological analysis and immunohistochemistry

BME domes with organoids were covered with 4% solution of formaldehyde (#252549, Sigma) for 10 minutes at room temperature. Then, BME domes were scraped off the plates, suspensions were transferred to conical tubes and centrifuged at 300 rcf for 5 minutes. Supernatants were removed and pellets were gently resuspended in hot HistoGel (#HG-4000-012, ThermoFisher). Histogel was solidified at room temperature, embedded in paraffin and sectioned. Tissue or organoid sections (3.5 μm) were de-paraffinized in tissue-clear (#1466, Sakura) and hydrated in ethanol solutions at decreasing concentrations before further treatment. H&E staining was performed using Mayer’s hematoxylin (5 minutes) and eosin (5 minutes), (01820 and 01650, Histolab Products AB). For immunohistochemical stainings, heat-induced antigen retrieval was performed at 95°C for 15 minutes in 10 mM sodium citrate, pH 6.0 (for Ki67 and CK7) or at 95°C for 20 minutes in Tris-EDTA buffer (10 mM Tris base, 1 mM EDTA, 0.05% Tween-20, pH 9.0) (for PAX8 and WT1). Endogenous peroxidase activity was blocked by incubation in 1% (v/v) hydrogen peroxide for 15 minutes. The sections were then incubated overnight in Shandon racks (Thermo Shandon) at 4°C with the primary antibodies (100 μL/section) diluted in Antibody Diluent (#S302283-2, Agilent) to the following concentrations; 1:1000 mAb PAX8 (#ab191870, Abcam); 1:100 mAb WT1 (#ab89901, Abcam); 1:500 mAb Ki67 (#275R-16, Cell Marque) and 1:100 mAb CK7 (#ab183344, Abcam). Envision horseradish peroxidase-labeled anti-rabbit IgG (K4003, Dako) was used for detection (100 μL/section) by incubating 45 minutes at room temperature, followed by development with NovaRED (Vector Laboratories) for 9 minutes as specified by the manufacturer. The sections were then counterstained with Mayer’s hematoxylin for 30 seconds. Both H&E and immunoperoxidase stained sections were finally dehydrated in ethanol solutions before mounting in tissue mount (#1467, Sakura).

Morphological features such as growth pattern and nuclear pleomorphism, in both tissue and organoid sections, were compared by two pathologists using a light microscope (Olympus BX46 Clinical Upright Microscope). Nuclear pleomorphism was scored as weak (1+), moderate (2+) or severe (3+). Similarly, the expression of each IHC marker was assessed in a blinded manner and independently by the two observers using light microscopy. After verifying the specificity of each immunostaining in the samples (nuclear stain for PAX8, WT1 and Ki-67; membranous and cytoplasmic for CK7), the staining intensity (negative = 0, weak = 1+, moderate = 2+, strong = 3+), the degree of homogeneity/variability of the immunostainings and the percentage of stained cells were recorded for each marker. Consensus on discrepant scores was obtained by simultaneous observation using a double-arm microscope.

### Whole-genome sequencing and analysis

Genomic DNA was extracted from snap-frozen organoids using DNAeasy Blood & Tissue kit (#69504, QIAGEN) and from snap-frozen tumor tissue or ascites cells, as well as from whole blood buffy coat (germline) samples using AllPrep DNA/RNA mini kit (#80204, QIAGEN). DNA quality was assessed using a Qubit fluorometer (Invitrogen). The samples were sequenced with Illumina HiSeq X Ten, BGISEQ-500 or MGISEQ-2000 at BGI Genomics Europe (Copenhagen, Denmark) and processed the Anduril 2 platform ^45^. Short-reads were aligned to human genome GRCh38.d1.vd1 using BWA-MEM ^46^ followed by duplicate removal with Picard Tools (http://broadinstitute.github.io/picard/) and base quality score recalibration with GATK ^47,48^.

Somatic mutations were called using GATK Mutect2 ^49^ with joint calling. A panel of normals generated from 105 HERCULES/DECIDER and 99 TCGA blood-derived normal samples was utilized. Mutations were annotated using CADD ^50^, ANNOVAR ^51^ and ClinVar ^52^. Non-silent or splicing variants pathogenic in ClinVar or non-benign with CADD phred score > 10 were considered functionally relevant. Mutation concordance was defined between two samples as Jaccard index using the number of shared and private mutations, where a mutation is present in a sample as long as at least one ALT read is observed.

Germline mutations used in CNV analysis come from a callset of 106 HERCULES/DECIDER normals using GATK ^53^. Variant quality score recalibration was allele-specific.

Mutational signature analysis was performed using COSMIC v3.1 signatures ^29,30^ adjusted for GRCh38 nucleotide frequencies. Mutations without VAF > 0.05 in any sample as well as those that only had ALT reads in one strand were excluded. Sample mutational profiles comprised mutations with at least one ALT read in that sample.

### CNV analysis

Copy-number segmentation was done with GATK ^47^. To collect the minor allele counts, all filtered biallelic (VAF 0.4-0.6) germline SNPs from each patient were used. Read-count collection used one-kilobase intervals. Both read and allelic count collection excluded regions listed in the ENCODE blacklist ^54^ and internal HERCULES/DECIDER blacklist, which includes regions that have abs(logR) > 0.2 in at least three of the 114 available normal samples.

After the segmentation, a reimplemented ASCAT algorithm ^55^ was used to estimate purity, ploidy, and allele-specific copy numbers. The original ASCAT R package was not directly applicable because it does not accept data segmented using external tools. Our implementation also uses VAF of homozygous TP53 mutations as additional evidence for the ploidy/purity estimates.

### Single-cell RNA Sequencing and analysis

Cryopreserved tissue samples were thawed and resuspended in culture medium immediately before sample processing for scRNA-seq and subsequently processed for scRNA-seq in the same way as dissociated organoid samples. Organoid cultures were dissociated in 2 mL of TrypLE Express for 25 minutes with occasional trituration by pipetting. The dissociated organoid cells were washed twice with 10 mL ice-cold PBS, and resuspended in 1 mL of PBS. The live cells were counted by Trypan blue exclusion method using Countess II counter (ThermoFisher). scRNA-seq libraries were prepared with Chromium Single Cell 3′ Reagent Kits (v.2.0 or v.3.1, 10× Genomics) and samples were sequenced on an Illumina NovaSeq 6000 instrument (Sequencing Unit of the Institute for Molecular Medicine Finland). The fastq files obtained from sequencing were processed using 10× Genomics Cell Ranger v5.0.0 for de-multiplexing, alignment, barcode processing and UMI quantification. GRCh38.d1.vd1 genome was used as reference and GENCODE v25 for gene annotation. Count matrices were loaded in Seurat (v3.2.2) ^56^ and initial filtering was applied to remove cells with <20% of mitochondrial reads. Next, uniform manifold approximation and projection (UMAP) was used for dimensionality reduction and initial clustering with k-means. At this stage, three major cell types were assigned based on the marker expression: epithelial tumor cells (WFDC2, PAX8, MUC16), stromal cells (COL1A2, FGFR1, DCN) and immune cells (CD79A, FCER1G, PTPRC). To ensure the quality of the cells used in subsequent analysis, further filtering was performed by removing stromal and immune cells with a number of UMI (in log scale) <11 and tumor cells with <12, resulting in total 2884, 8070 and 8572 cells of particular type, respectively.

Patient-specific molecular markers for tumor samples and their derived organoids were extracted using a logistic regression. The top patient-specific tumor markers with a logFC >2 were plotted and compared against organoid patient-specific markers.

CNV profiles for the individual cancer cells were obtained with InferCNV of the Trinity CTAT Project ^34^ using the stromal and immune cells as reference. Next, Leiden algorithm was used for the determination of the underlying subclonal populations based on the CNV profiles. The similarity matrix for the subclones was calculated by measuring the cosine distance between the consensus CNV status of each subclone.

### Drug response profiling

Fully grown organoids were extracted from BME-2 domes as described above. Cell pellets were resuspended in BME-2 and 10 μL of the suspension was plated into each well (omitting outside border wells) of pre-cooled 384-well Ultra-Low Attachment microplates. Seeding density was adjusted to the expansion rate of particular organoid culture using the following formula: Number of organoid culture droplets seeded into a single microplate = culture passaging ratio x 30. Gel droplets were solidified in a humidified cell culture incubator at 37°C for 30-45 minutes and covered with 40 μL of culture-appropriate growth medium. Culturing medium was exchanged every 2-3 days using EL406 plate washer (BioTek). 2-3 days before drug addition, in half of the plates, M1/M2 medium was exchanged to HPLM (#A4899101, ThermoFisher), supplemented with relevant niche factors from M1/M2, omitting GlutaMax, nicotinamide and N-acetyl-cysteine, which majorly impact cell metabolism. After 7-10 days of growth post-seeding (depending on the growth rate of each organoid culture), experimental drugs, DMSO (vehicle control) or 10 μM bortezomib (positive control) were added to the microplates using Echo 550 Liquid Handler (Labcyte). Experimental compounds were added in five 10-fold dilutions (carboplatin at 10 nM-100 μM, other compounds at 1 nM-10 μM). Microplates were incubated at 37°C for 96 hours. Subsequently, 25 μL of the culture medium was exchanged to staining solution, containing CellTox Green reagent (#G8743, Promega, final concentration 1X) and Hoechst 33342 (#B2261, Sigma, final concentration 5 μg/mL). Plates were incubated for 2 hours at room temperature and imaged in an ImageXpress Micro Confocal automated fluorescence microscope (Molecular Devices). A single image per well for each fluorophore was taken using 10X objective. Survival indices were estimated by image analysis using MetaXpress (Molecular Devices) software. Nuclei were identified as individual objects based on Hoechst 33342 staining and computationally enlarged. Objects which overlapped with CellTox Green signal were counted as dead cells and negative as viable cells. Data points coming from microplate wells that lost the BME gels with embedded organoids during plate processing (for example, during media exchanges) were removed from the analysis. Fraction of viable cells was estimated in each well. Survival index in each well was estimated by normalization to negative (100% viability) and positive (0%) controls. Dose-response curves were generated using non-linear log(inhibitor) vs response (three parameters) fit and AUC calculated in GraphPad Prism. For AUC calculation for combination of carboplatin and 0.1 μM paclitaxel, only three highest concentrations of carboplatin were used (1-100 μM) in order to avoid exaggerated effect of 0.1 μM paclitaxel on overall drug combination effect at low carboplatin concentrations (1-100 nM). To simplify comparisons between different samples and medium conditions, AUCs were transformed for particular treatments using Z-score normalization. For correlation of drug response to patient outcomes, normalized CA125 change was calculated as a difference in CA125 level before chemotherapeutic treatment and the final, post-treatment CA125 level in that patient (the last data point in the yellow area in patient CA125 graphs over time) before or at the start of other treatment, expressed as a percentage of the maximal, patient-specific CA125 blood level in the relevant period.

### Statistical analyses

Mean organoid radius values were compared with unpaired two-tailed *t* test using GraphPad Prism. Correlations between single-cell expression levels of organoid markers and tumor markers were calculated using Pearson’s *R*, assuming linear relationship. Correlations between normalized blood CA125 level change in patients and organoid-based normalized AUC Z-score were not assumed to be linear and were calculated using Spearman’s r. *P* values <0.05 were considered significant.

## Supporting information

Supplemental figures S1-S6; Step-by-step protocols for organoid culture

## ACKNOWLEDGEMENTS

We thank Professor Kim Jensen for kindly providing R-Spondin 1- and Wnt-conditioned media for this study. We gratefully acknowledge the administrative support of Tiia M. Pelkonen. We thank Kylie Gallagher of Massachusetts Institute of Technology for assistance with medium optimization experiments. The results here are in part based upon data generated by the TCGA Research Network: https://www.cancer.gov/tcga. Figures 1a, 3b and 6a were created with BioRender. This work was supported by the European Union’s Horizon 2020 research and innovation programme under Grant Agreements no. 667403 for HERCULES (O.C., K. H., J.Hy.,S.H., A. Vä, K.W.), no. 965193 for DECIDER (A.Vi., O.C., J.Hy., S.H) and no. 845045 for RESIST3D (W.S.); Danish Cancer Society, grant no. R204-A12322 (W.S.); Novo Nordisk Foundation: Novo Nordisk Foundation Center for Stem Cell Biology, grant no NNF17CC0027852 (K.W.), High Content CRISPR Screens (HCCS) facility, grant number NNF-0061734 (K.W.); Innovation Fund Denmark and Academy of Finland for ERA PerMed JTC2020 PARIS project, grant numbers 0204-00005B (K.W.,) and 344697 (A.Vä.), respectively; Academy of Finland, grant numbers 319243 (A. Vä) and 1289059 (A.Vä); The Sigrid Jusélius Foundation (A.Vä); The Cancer Foundation Finland (A.Vä).

## AUTHOR CONTRIBUTIONS

Conceptualization: W.S., K.W. Patient recruitment and management: O.C., J.Hy. Patient sample processing:T.L., K.K., J.Hu., A.Vi, K.H. Organoid culture: W.S., L.G., D.B. Histological staining: A.Vi, M.C.K., M.K.G.H.,I.M.L. Pathologic analysis: P.C., E.S. Genomic analysis: Y.L., K.L., J.O. Single-cell transcriptomics and analysis: M.M.F., E.P.E., A.Vä. Drug response experiments and analysis: W.S., E.J.P. K.V., Y.C. Supervision: L.E., K.H., O.C., J.Hy., S.H., A.Vä., K.W. Resources and funding acquisition: W.S., D.B., L.E., K.H., O.C.,J.Hy., S.H., A.Vä., K.W. Writing—original draft: W.S. Writing—review and editing: W.S., L.G., M.M.F., Y.L,K.L., M.C.K., J.O., D.B., E.J.P., K.V., Y.C., E.P.E., M.K.G.H., I.M.L., T.L., K.K., J.Hu., A.Vi., L.E., P.C., E.S., K.H., O.C., S.H., A.Vä., K.W.

## Declaration of interests

The authors declare no competing interests.

**Materials & correspondence:** Organoid cultures EOC41_pOme, EOC733_pPer and EOC733_iOme and associated clinical data will be available upon publication of this work in a peer-reviewed journal to the research community through the Auria Biobank (https://www.auria.fi/biopankki/en/), based on patients’ informed consents. Genomic data, including mutation, copy-number and signature profiles are available through an interactive visualization in GenomeSpy, accessible at: https://csbi.ltdk.helsinki.fi/p/senkowski_et_al_2022/. Genomic sequence data will be available upon publication at the European Genome-phenome Archive (EGA), which is hosted by the EBI and the CRG, under accession number EGAS00001004714. Transcriptomic sequence data will be available from EGA upon publication. For inquiries regarding the availability of other materials, contact K.W. For technical details or inquiries regarding the organoid culture protocol, contact W.S.

