## Supplemental figures S1-S6; Step-by-step protocols for organoid culture for "A platform for efficient establishment, expansion and drug response profiling of high-grade serous ovarian cancer organoids"

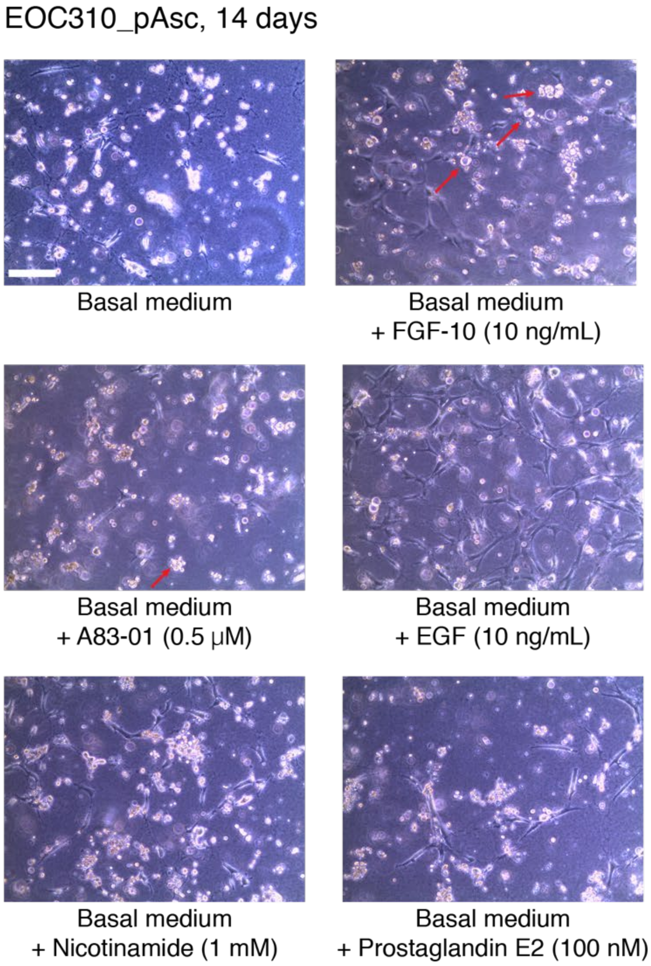

**Figure S1. Influence of individual additives on short-term HGSC organoid formation. Related to Figure 1.**
Phase-contrast images of EOC310_pAsc cells, embedded in BME and cultured for 14 days in the Basal Medium, supplemented with individual additives, as indicated. Red arrows indicate formation of coherent, three-dimensional multicellular clusters. 10X magnification; scale bar, 100 μm.

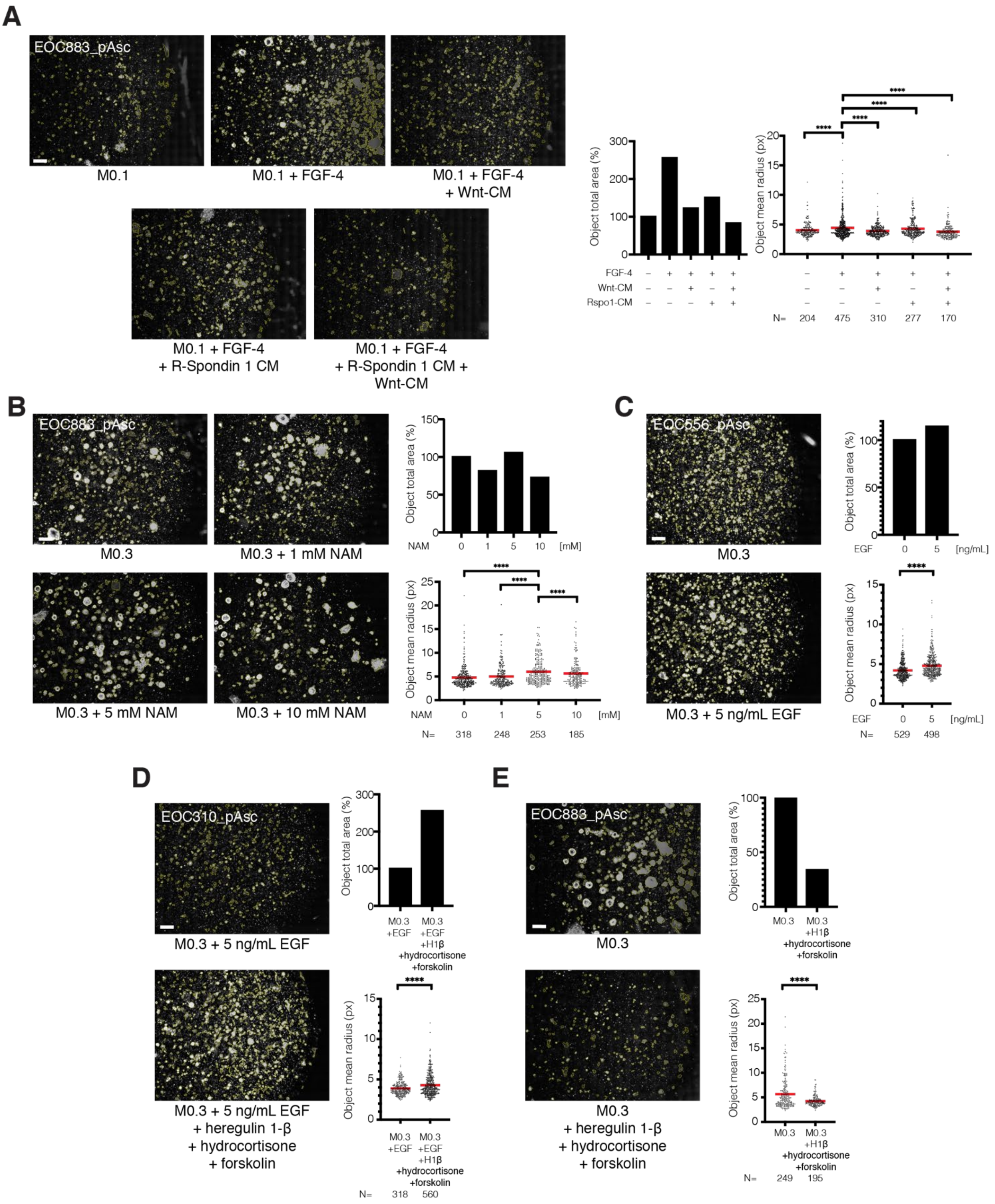

**Figure S2. Establishment of new HGSC organoid media formulations. Related to Figure 1.**
(A) *Left:* Phase-contrast images of basement membrane extract (BME) droplets with objects (outlined in yellow) identified with CellProfiler. EOC883_pAsc cells were cultured in M0.1 or M0.1 supplemented with FGF-4 (10 ng/mL), Wnt conditioned medium (Wnt-CM, 50% v/v) and/or R-Spondin 1 conditioned medium (R-Spondin 1 CM, 25% v/v) for 38 days (passaged once on day 17). Scale bar, 200 µm. *Right:* Total area of objects and mean (marked with a line) object radius in the particular picture, estimated using CellProfiler. (B) *Left:* Phase-contrast images of BME droplets with objects identified as above. EOC883_pAsc cells were cultured in M0.3 or M0.3 supplemented with nicotinamide (NAM, 1, 5 or 10 mM) for 38 days (passaged once on day 19). Scale bar, 200 µm. *Right:* Mean object radius in the particular picture, as above. (C) *Left:* Phase-contrast images of BME droplets with objects identified as above. EOC556_pAsc cells were cultured in M0.3 or M0.3 supplemented with EGF (5 ng/mL) for 33 days (passaged once on day 17). Scale bar, 200 µm. *Right:* Mean object radius in the particular picture, as above. (D, E) *Left:* Phase-contrast images of BME droplets with objects identified as above. EOC310_pAsc (D) or EOC883_pAsc (E) cells were cultured in M0.3 supplemented with 5 ng/mL EGF (D) or M0.3 (E) or these formulations supplemented with 37.5 ng/mL heregulin-1β, 0.5 µg/mL hydrocortisone and 5 µM forskolin for 35 days (passaged once on day 19, (D)) or 39 days (passaged once on day 19, (E)). Scale bar, 200 µm. *Right:* Mean object radius in the particular picture, as above.
**** = p<0.0001, unpaired two-tailed t-test.

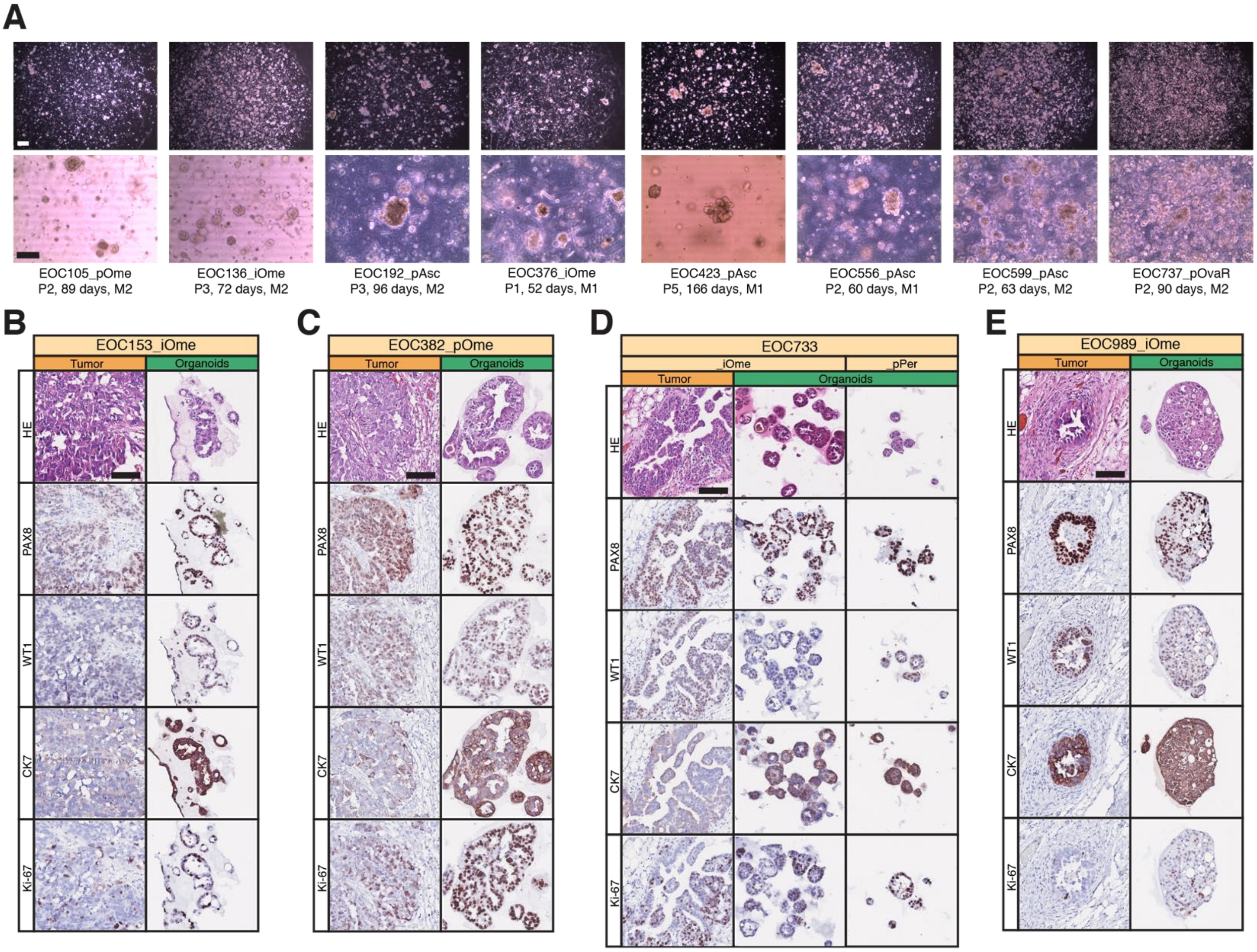

F

| Patient | EOC153 | | EOC382 | | EOC733 | | | EOC989 | | EOC1120 | | |
|---|---|---|---|---|---|---|---|---|---|---|---|---|
| Sample | _iOme | | _pOme | | _iOme | | _pPer | _iOme | | _pOme | | _rAsc |
| | Tissue | Organoids | Tissue | Organoids | Tissue | Organoids | Organoids | Tissue | Organoids | Tissue | Organoids | Organoids |
| Nuclear pleomorphism | 3+ | 1+ | 3+ | 3+ | 3+ | 3+ | 3+ | 3+ | 3+ | 3+ | 3+ | 3+ |
| PAX8 | Variable (0 to 3+) | 3+ | Variable (0 to 3+) | 3+ | Variable (0 to 3+) | 3+ | 3+ | 3+ | 3+ | Variable (0 to 3+) | 3+ | 3+ |
| WT1 | Variable (0 to 3+) | Variable (2+ to 3+) | Variable (0 to 3+) | Variable (2+ to 3+) | Variable (0 to 3+) | Variable (0 to 3+) | Variable (0 to 3+) | 3+ | Variable (0 to 3+) | Variable (1+ to 3+) | Variable (0 to 3+) | 3+ |
| CK7 | Variable (0 to 3+) | 3+ | Variable (0 to 3+) | 3+ | Variable (0 to 3+) | 3+ | 3+ | 3+ | 3+ | Variable (1+ to 3+) | 3+ | 3+ |
| Ki-67 | Variable (0 to 3+) | Variable (2+ to 3+) | Variable (0 to 3+) | 3+ | Variable (2+ to 3+) | 3+ | 3+ | 3+ | 3+ | Variable (0 to 2+) | Variable (1+ to 3+) | Variable (1+ to 3+) |

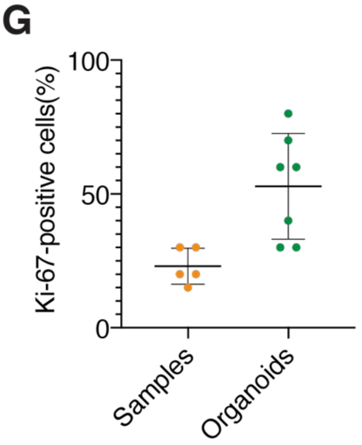

**Figure S3. Immunohistochemical comparison of organoid models and tissues of origin. Related to Figure 3.** (A) Brightfield/phase-contrast images of failed cultures depicting initial 3D structure formation and cellular growth. Scale bars, 200 µm (*top*) and 100 µm (*bottom*). (B-E) HE and IHC stainings (for indicated markers) of EOC153_iOme, EOC382_pOme, EOC733_iOme and EOC989_iOme tumor tissues and matching organoids. Additionally, EOC733_pPer organoids were stained (C). Scale bar, 100 µm. (F) Pathological assessment and scoring of the stained tissues. Organoids demonstrate morphological features similar to the original tissue, including nuclear pleomorphism, adenopapillary growth pattern and positive staining for PAX8, WT1 and CK7. They are also more proliferative than the original tissue, depicted by higher Ki-67 expression. (G) Comparison of estimated cancer cell Ki-67 positivity between organoids and original tissues, based on Ki-67 IHC staining, presented as mean ± s.d.

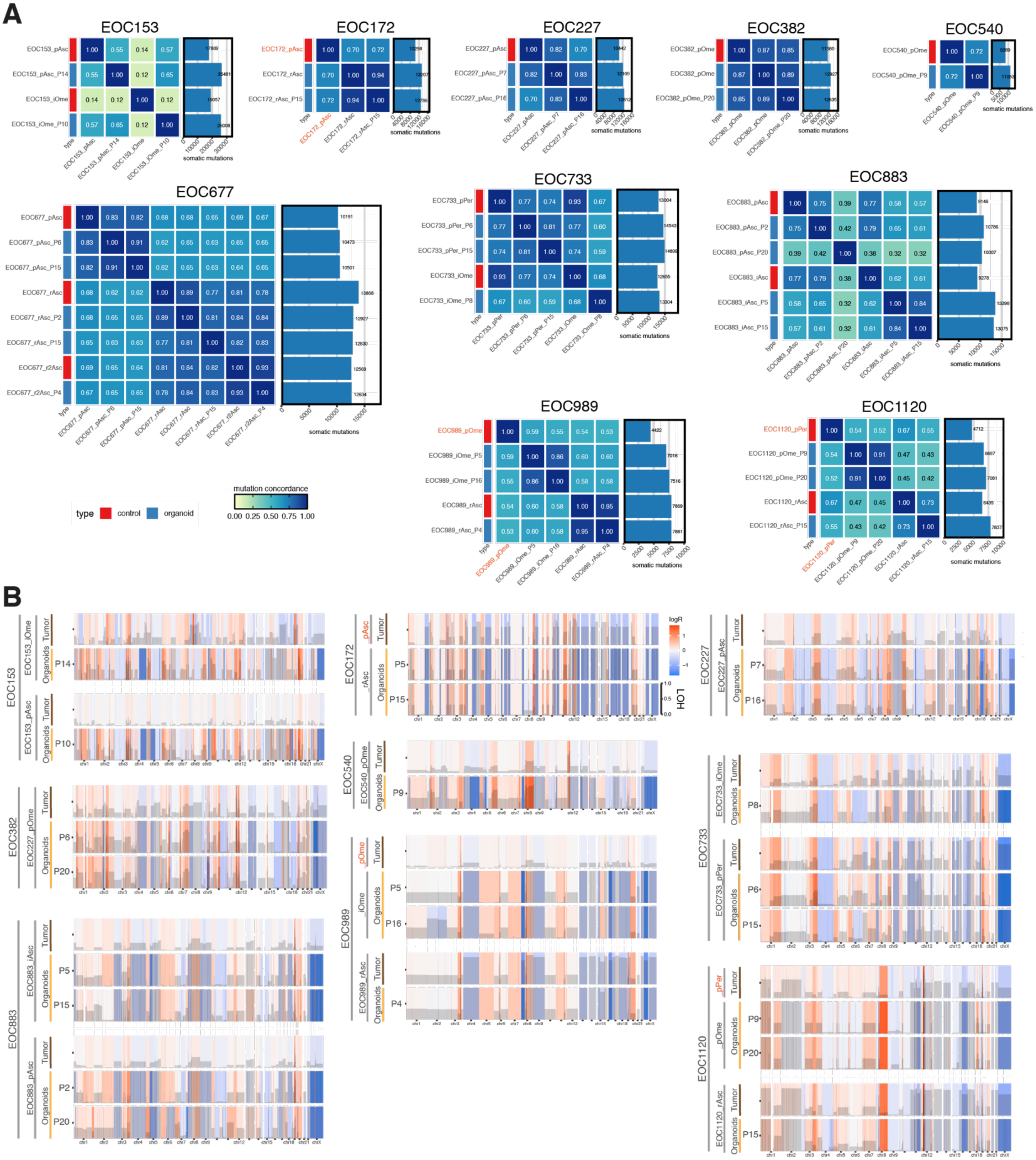

**Figure S4. Comparison of genomic landscapes of organoids and original tumor tissues. Related to Figure 4.**
(A) Total number of detected somatic mutations indicated for each sample. Sample names are typed in orange, where tumor tissue from a different metastatic location/clinical progression stage than the one used for organoid derivation is presented (due to limited matching tissue availability for sequencing). (B) Genome-wide CNV analysis of tumor tissue and corresponding organoids. Copy number changes are expressed as logR and color-coded. The extent of LOH is displayed with grey bars. Passage numbers (P) at sequencing are indicated for organoid cultures. Sample names are typed in orange, where tumor tissue from a different metastatic location/clinical progression stage than the one used for organoid derivation is presented (due to limited matching tissue availability for sequencing).

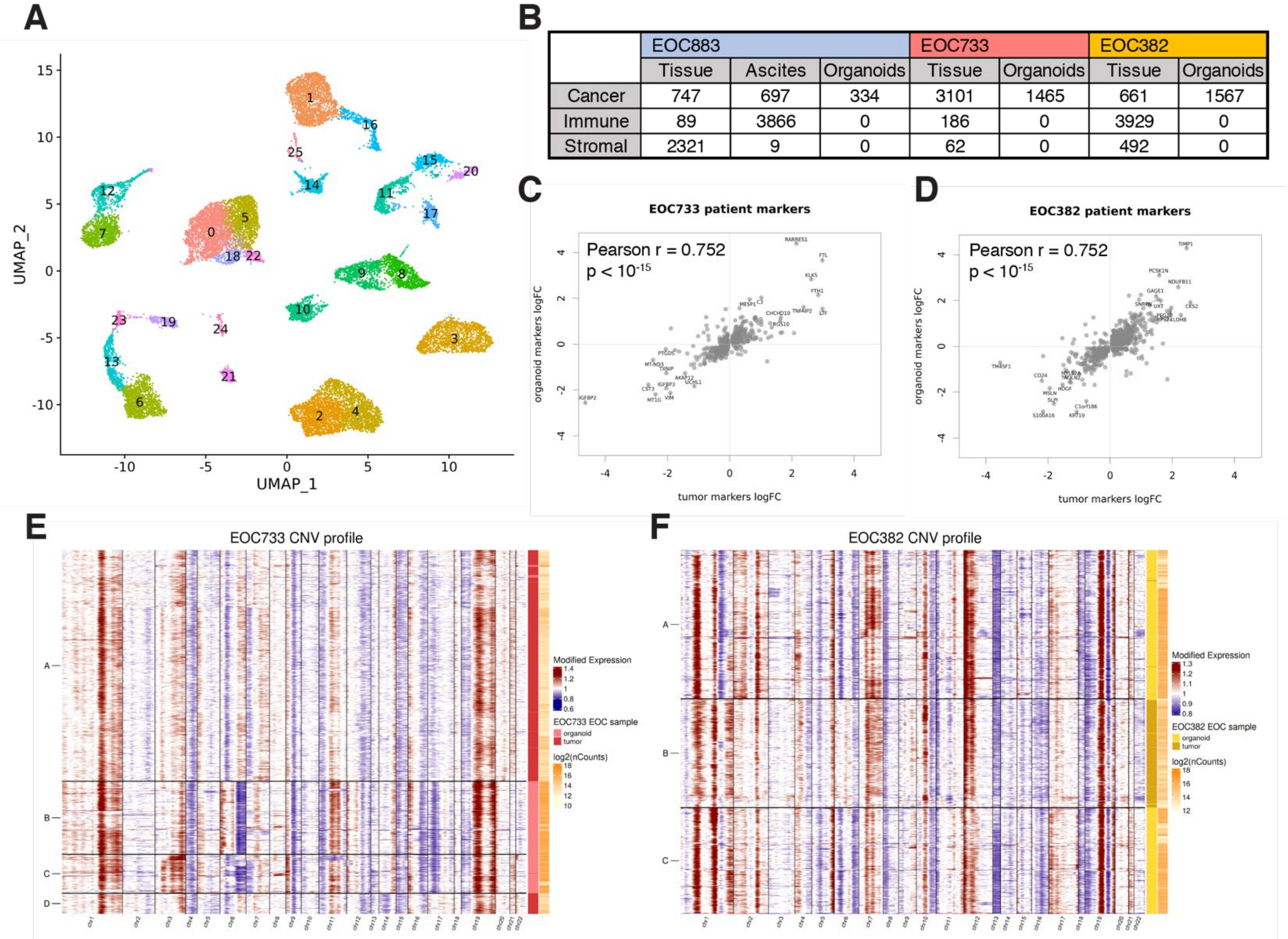

**Figure S5. scRNA-seq characterization of HGSC organoids. Related to Figure 5.**
(A) UMAP visualization of 19,526 cells from tumor samples: EOC883_pAsc, EOC883_pAdn, EOC382_pOme and EOC733_pPer and corresponding organoids (except for EOC883_pAdn), assigned to 26 subclusters (indicated by different colors and numbers) through unsupervised clustering. (B) Number of cells in analyzed samples assigned to a particular cell type (cancer, stromal or immune). (C, D) Pearson correlation plots of patient-specific markers expression in EOC733_pPer (C) or EOC382_pOme (D) tumor samples and corresponding organoids. (E, F) Single-cell CNV plots from EOC773_pPer (E) or EOC382_pOme (F) tumors and organoids, inferred using InferCNV and classified into 4 subclusters.

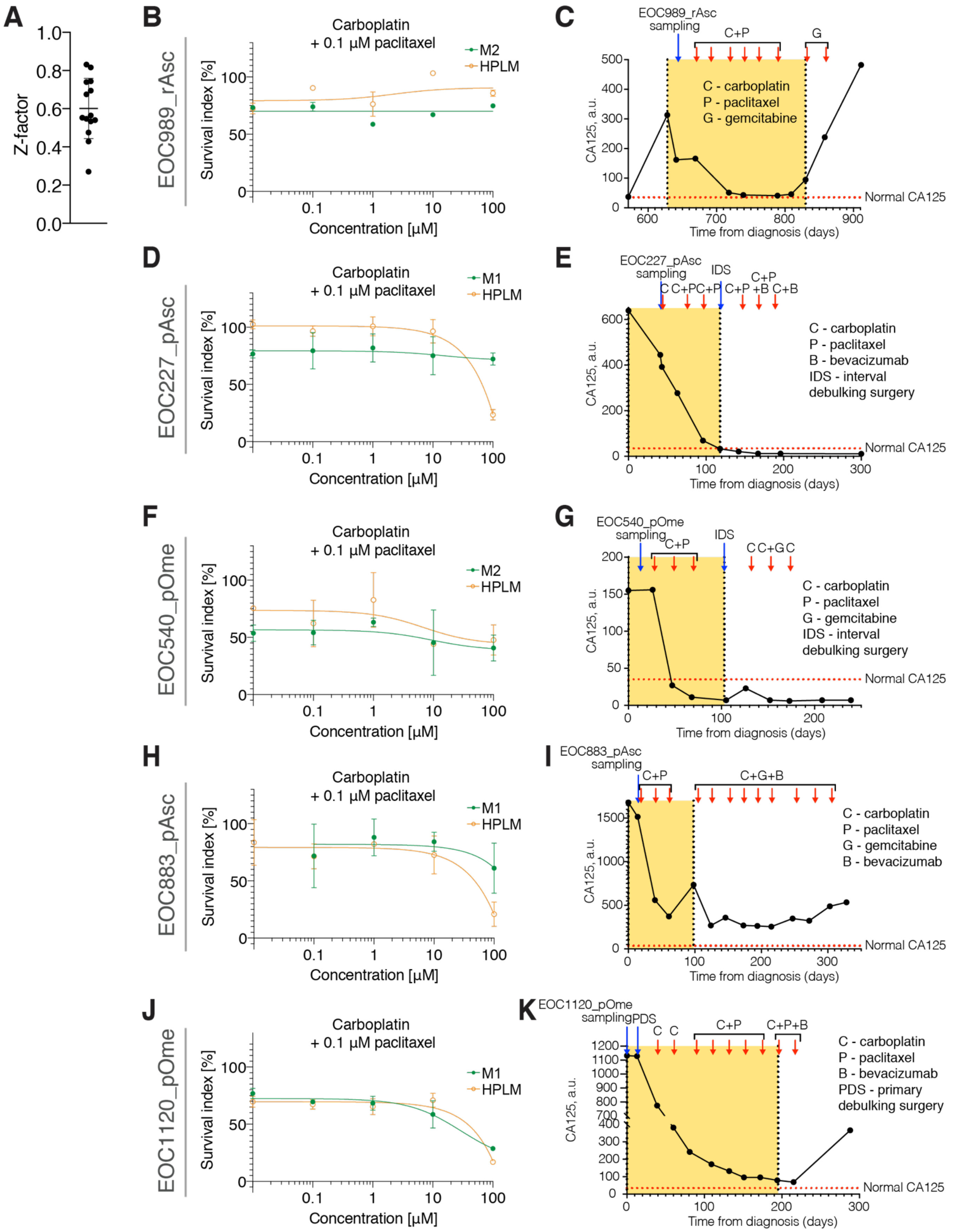

**Figure S6. HGSC organoid drug responses and patient clinical outcomes. Related to Figure 6**
(A) Z-factors in the drug response profiling experimental 384-well microplates. Presented as mean ± s.d. (n=14). (B, D, F, H, J) Dose-response curves of EOC989_rAsc (B), EOC227_pAsc (D), EOC540_pOme (F), EOC883_pAsc (H) or EOC1120_pOme (J) organoids treated with carboplatin at indicated concentrations + 0.1 µM paclitaxel, in M1/M2 or HPLM. Results are shown as mean ± s.d. (n = 1-3). (C, E, G, I, K) CA125 blood levels of patients EOC898 (C), EOC227 (E), EOC540 (G), EOC883 (I) or EOC1120 (K) over time, during first-line therapy (E, G, I, K) or at relapse (C). Period relevant for comparison with *in vitro* drug response indicated with yellow rectangles. Normal CA125 range (<35 a.u.) indicated with red dotted lines.

| Patient no. | Patient code | FIGO stage at diagnosis | Sample name | Tumor deposit source | Clinical progression stage at sampling | Sample tumor purity | Successful organoid derivation | Organoid tumor purity (latest available passage) | Time in culture to reach stable expansion (days) | Successful resuscitation from frozen organoids |
|---|---|---|---|---|---|---|---|---|---|---|
| Samples used for organoid derivation and medium optimization | | | | | | | | | | |
| 1 | EOC105 | IIIC | EOC105_pOme | Omentum | Primary | Unknown | No | | | |
| 2 | EOC136 | IVA | EOC136_pAsc | Ascites | Primary | 12% | No | | | |
| | | | EOC136_iOme | Omentum | Interval | 72.9% | No | | | |
| 3 | EOC153 | IVA | EOC153_pAsc | Ascites | Primary | 15.5% | Yes | 100% | 120 | Yes |
| | | | EOC153_iOme | Omentum | Primary | 70.5% | Yes | 99% | 185 | Yes |
| 4 | EOC172 | IVA | EOC172_pOme | Omentum | Primary | 40.5% | No | | | |
| | | | EOC172_rAsc | Ascites | Recurrence | 0%* | Yes | 100% | 100 | Yes |
| 5 | EOC192 | IIIC | EOC192_pAsc | Ascites | Primary | 35.5% | No | | | |
| | | | EOC192_pOvaL | Left ovary | Primary | Unknown | No | | | |
| 6 | EOC227 | IVA | EOC227_pAsc | Ascites | Primary | 35.9% | Yes | 98.50% | 102 | Yes |
| 7 | EOC376 | IIIC | EOC376_iOme | Omentum | Interval | Unknown | No | | | |
| 8 | EOC382 | IIIC | EOC382_pOme | Omentum | Primary | 35% | Yes | 97.5% | 30 | Yes |
| 9 | EOC41 | IIIC | EOC41_pOme | Omentum | Primary | 45.5% | Yes | Testing ongoing | 328 | Yes |
| 10 | EOC423 | IIIC | EOC423_pAsc | Ascites | Primary | Unknown | No | | | |
| | | | EOC423_pOme | Omentum | Primary | 70.2% | No | | | |
| | | | EOC423_iOvaR | Right ovary | Interval | 100% | No | | | |
| 11 | EOC473 | IVB | EOC473_pAdn | Adnex | Primary | 72% | No | | | |
| | | | EOC473_iPer | Peritoneum | Interval | Unknown | No | | | |
| 12 | EOC540 | IIIC | EOC540_pOme | Omentum | Primary | 22% | Yes | 99.50% | 120 | Yes |
| 13 | EOC556 | IIIC | EOC556_pAsc | Ascites | Primary | 17.8% | No | | | |
| | | | EOC556_iBow | Bowel | Interval | 8.5% | No | | | |
| 14 | EOC599 | IVA | EOC599_pAsc | Ascites | Primary | 18% | No | | | |
| | | | EOC599_iOme | Omentum | Interval | 10% | No | | | |
| 15 | EOC677 | IIIC | EOC677_pAsc | Ascites | Primary | 45.4% | Yes | 100% | 86 | Yes |
| | | | EOC677_rAsc | Ascites | Recurrence | 40.8% | Yes | 97.5% | 26 | Yes |
| | | | EOC677_r2Asc | Ascites | 2nd recurrence | 67.5% | Yes | 100% | 49 | Yes |
| 16 | EOC737 | IIIC | EOC737_pOvaR | Right ovary | Primary | 50.7% | No | | | |
| 17 | EOC733 | IVA | EOC733_pPer | Peritoneum | Primary | 97% | Yes | 96.5% | 91 | Yes |
| | | | EOC733_iOme | Omentum | Interval | 51.5% | Yes | 100% | 140 | Yes |
| 18 | EOC883 | IIIC | EOC883_pAsc | Ascites | Primary | 18.5% | Yes | 100% | 35 | Yes |
| | | | EOC883_iAsc | Ascites | Interval | 27.9% | Yes | 99.00% | 85 | Yes |
| 19 | EOC989 | IVA | EOC989_iOme | Omentum | Interval | 5% | Yes | 100% | 55 | Yes |
| | | | EOC989_rAsc | Ascites | Recurrence | 90.8% | Yes | 99.50% | 49 | Yes |
| 20 | EOC1120 | IVB | EOC1120_pOme | Omentum | Primary | Unknown | Yes | 100% | 177 | Yes |
| | | | EOC1120_rAsc | Ascites | Recurrence | 80% | Yes | 99% | 70 | Yes |
| Samples used only for medium optimization | | | | | | | | | | |
| 21 | EOC310 | | EOC310_pAsc | Ascites | Primary | Unknown | N/A | | | |

**Table S1. Overview of samples used in the study. Related to Figures 1-4**

*Cancer cells not detectable in the sample using WGS

| Media supplements tested for organoid establishment and long-term culture | | | |
|---|---|---|---|
| **Growth factors** | **Manufacturer, product no.** | **Concentrations tested** | **Effect on organoid derivation** |
| FGF-2 | Peprotech, #100-18B | 10 ng/mL | Harmful |
| FGF-4 | Peprotech, #100-31 | 10 ng/mL | Beneficial |
| FGF-7 | Peprotech, #100-19 | 10 ng/mL | Neutral |
| FGF-10 | Peprotech, #100-26 | 10 ng/mL | Beneficial |
| EGF | Peprotech, #AF-100-15 | 5 ng/mL | Harmful or beneficial |
| | | 10 ng/mL | Harmful |
| | | 50 ng/mL | Harmful |
| IGF-I | Peprotech, #100-11 | 20 ng/mL | Neutral |
| | | 100 ng/mL | Neutral |
| VEGF | Peprotech, #AF-100-20 | 10 ng/mL | Neutral |
| **Other proteins** | | | |
| Heregulin-1β | Peprotech, #100-03 | 5 nM | Harmful or beneficial |
| BMP-2 | Thermo Fisher, #PHC7145 | 10 ng/mL | Neutral |
| Jag-1 | | 1 μM | Neutral |
| R-Spondin 1 | Peprotech, #120-38 | 100 ng/mL | Neutral |
| | | 400 ng/mL | Neutral |
| | | 1 μg/mL | Harmful |
| R-Spondin 3 | Peprotech, #3500-RS-025 | 250 ng/mL | Neutral |
| Noggin | Peprotech, #120-10C | 100 ng/mL | Harmful |
| **Hormones** | | | |
| β-estradiol | Sigma, #E8875 | 100 nM | Beneficial |
| Hydrocortisone | Sigma, #H0888 | 100 ng/mL | Harmful or beneficial |
| | | 500 ng/mL | Harmful or beneficial |
| Follicle-stimulating hormone | R&D Systems, #5925-FS-010 | 10 ng/mL | Neutral |
| | | 50 ng/mL | Neutral |
| Gonadotropin-stimulating hormone | Sigma, #L8008 | 10 ng/mL | Neutral |
| | | 50 ng/mL | Neutral |
| Triiodothyronine | Sigma, #T6397 | 0.1 ng/mL | Harmful |
| | | 1 ng/mL | Harmful |
| | | 10 ng/mL | Harmful |
| Prostaglandin E2 | MedChemExpress | 10 nM | Neutral |
| | #HY-101952 | 1 mM | Harmful |
| **Small-molecule inhibitors** | | | |
| A83-01 | Sigma, #SML0788 | 0.5 μM | Beneficial |
| SB202190 | MedChemExpress, #HY-10295 | 0.5 μM | Beneficial |
| | | 3 μM | Beneficial |
| | | 10 μM | Beneficial |
| CHIR-99021 | MedChemExpress, #HY-10182 | 2.5 μM | Harmful |
| Idasanutlin | MedChemExpress, #HY-15676 | 0.1 μM | Harmful |
| Forskolin | MedChemExpress, #HY-15371 | 5 μM | Harmful or beneficial |
| | | 10 μM | Harmful or beneficial |
| **Conditioned media** | | | |
| Rspo1-conditioned medium | Gift from prof. Kim Jensen | 25% v/v | Harmful |
| Wnt-conditioned medium | Gift from prof. Kim Jensen | 20% v/v | Harmful |
| | | 50% v/v | Harmful |
| **Other** | | | |
| Nicotinamide | Sigma, #N0636 | 1 mM | Beneficial |
| | | 5 mM | Beneficial |
| | | 10 mM | Beneficial |

**Table S2. Overview of tested media additives and their effects on HGSC organoid culture. Related to Figures 1-2.**

| Data for success rate calculation in previous studies come from: | |
|---|---|
| Maenhoudt et al. (2020) | Table 1 |
| Hoffmann et al. (2020) | Table EV3 |
| Kopper et al. (2019) | Extended Data Fig. 2a and Supplementary Table 4 |

**Table S3. Data sources for the calculation of long-term organoid culture success rate in previous studies. Related to Figure 2.**

### PROTOCOLS FOR ORGANOID CULTURE

*High-Grade Serous Ovarian Cancer Organoids*

***Patient sample:** count cells.
**Frozen organoids:** don't

### Starting a culture from cryopreserved material (Patient Samples and Organoids)

#### Materials

- Dry ice and a Styrofoam box.
- Culture media (only M1).
- Y-27632 stock solution
- BME Type 2 (#3533-010-02, R&D Systems)
- 15-mL polypropylene snap cap Falcon tubes (1 per sample).
- 50-mL Falcon tubes.
- Pipette set and tips.
- Water bath set to 37ºC.
- 6-well culture plates Nunc Cell-Culture Treated (ThermoFisher: 140685).
- Multistep electronic pipette and tips.
- Cold, sterile 1X PBS [-] $CaCl_2$ [-] $MgCl_2$.
- M1 and M2.
- 10-mL serological pipettes.

#### Instructions (this section covers how to thaw frozen material)

##### *Before starting the protocol*

1. Fill a Styrofoam box with dry ice for short-term patient samples storage.
2. Place 6-well cell culture plates in the incubator (they need to be pre-heated before pipetting the BME-2 to allow instant gel polymerization).
3. Turn on the water bath (37ºC).

##### *Beginning of the protocol*

4. Aliquot 20 mL per sample of M1 in 50-mL Falcon tubes.
5. Warm up the media at 37ºC by placing the Falcon tubes in the water bath for 10-15'.
6. Transfer 10 mL of <u>M1 (without Y-27632)</u> to the snap cap Falcon tubes.
7. Defrost the samples by placing the cryovials in the water bath for 1-2'.
   a. Remember to swirl the cryovials so that the heat distributes homogeneously.
8. Add around 1 mL of M1 to each cryovial, mix 2-3 times, and transfer the content to the snap cap Falcon tubes. Rinse the cryovial with extra media.
9. Spin down the samples (200 or 300g, 5').
   a. 200g for patient samples.
   b. 300g for organoids.
10. Gently aspirate the supernatant.
    a. Be <u>very careful</u> because the pellet is usually quite loose.
11. Prepare 10 mL <u>M1 + Y-27632 (Fc: 5 µM)</u> per sample.
12. Transfer <u>10 mL of M1 + Y-27632</u> to the tube with the pellet and re-suspend it by pipetting around 10 times with the P1000.
    a. Remember not to over-pipette the cells too much when re-suspending. Having some cell clusters is beneficial for culturing.
13. If thawing a <u>patient sample</u>, count the cells. If thawing organoids, don't count them (just seed according to the split ratio).
14. Spin down the sample (200 or 300G, 5').
15. Gently aspirate the supernatant.
16. Go to Sample Seeding.

##### *Gel preparation (performed simultaneously with the procedure above)*

1. Slowly thaw the BME-2 (recommended to be performed on ice, as the gel polymerizes at room temperature).
   a. Remember to occasionally swirl the vial and place it in ice when defrosted. Never mix it by inversion.

2. Place 2 15-mL snap cap Falcon tubes in the ice bucket – one empty and the other with 1-2 mL of sterile PBS.
3. When the gel is defrosted, dilute it with **cold, sterile PBS**. Avoid introducing air bubbles. This step must be performed early so that any air bubbles have time reach the surface of the gel and disappear.
    a. Take the BME vial and gently re-suspend the content with the P1000.
    b. Transfer a desired amount of BME to a snap cap Falcon tube **together with the pipette tip** (since it contains a lot of product).
    c. Add cold, sterile PBS to obtain final protein BME concentration of 7.5 mg/mL and mix it until obtaining a homogenous solution.
    d. Remember to prep ≈ 100-200 µL of extra gel (as BME is a viscous solution, the volume indicated by the pipette is not exact and you will need some dead volume)
    e. each gel batch has a different protein concentration. We try to work with a protein concentration of around 7.5 mg/mL. The ideal gel concentration is 7.5-8 mg/mL, and the minimum required is 7 mg/mL. Thus, every batch is diluted differently (10-15% of PBS v/v).

*Sample seeding*

4. Gently mix the BME solution with the P1000. Avoid introducing air bubbles.
5. Take out 1 plate from the incubator and describe it with patient ID, sample, passage number, medium, and date.
6. Transfer the desired amount of the gel solution (200 µL + 50 µL extra volume per 10 droplets in a single well of 6-well plate) to the cell pellet and gently re-suspend until obtaining a homogenous solution (pipette between 10-14 times).
    a. **Patient samples:** re-suspend to obtain a density of min $10^6$ live cells/mL of BME-2.
    b. **Organoids:** follow the ratio on the tube.
7. Seed the cells with the electronic or manual pipette. **Seed 10 droplets of gel per plate (20 µL/droplet)**.
    a. Remember to place the droplets far enough from each other and from the walls to avoid merging (see Organoid Passaging protocol).
    b. **TIP:** decrease the aspiration speed to avoid air bubbles.
8. If seeding multiple samples, mix the gel between them.
9. Place the plates in the incubator for 45' to solidify the BME.

*During the 45-minute break*

10. Aliquot the desired amount (3 mL/well) of M1 and M2 media in Falcon tubes.
11. Add Y-27632 ($F_c$ 5µM) and mix.
12. Warm up the media at approx. 37ºC by placing the Falcon tubes in the water bath for approx. 10-15'.

*After the 45 minute break*

13. Take out the plates from the incubator and check the gels.
14. Finally, gently add 3 mL of media to each well (M1/M2 + Y-27632) with a **10-mL pipette.**
    a. Fresh patient samples should be cultured in M1 and M2 in parallel in order to determine (over a few passages) which medium formulation is preferred by the particular sample.
15. Place the plates in the incubator.

### Medium Change (6-well plates)

**The media are exchanged 3 times per week, every 2-3 days (usually Mondays, Wednesdays and Fridays).**

#### Materials

- Culture media (M1 and M2).
- 1X Sterile PBS [-] $CaCl_2$ [-] $MgCl_2$.
- 50-mL Falcon tubes.
- Serologic pipettes.
- Sterile, Pasteur glass pipettes for the vacuum pump.

#### Instructions

1. **Keep the media in the fridge while not using them.**
2. Turn on the water bath (37ºC).
3. In the meantime, take out the plates from the incubator and examine the cells. Calculate the amount of media and PBS that will be required.
    a. Each well must contain 3 mL of fresh media.
    b. For the washes, approximately 1 mL of PBS per well is required.
4. Aliquot the desired amount of M1, M2, and sterile PBS in Falcon tubes, and **place the media bottles back in the fridge**.
5. Warm up the media and the sterile PBS at approx. 37ºC in the water bath for approx. 10-15’.
    a. The media and PBS must be warm to prevent the gels from depolymerizing.
6. Take out the plates from the incubator.
7. Gently aspirate the old media.
    a. To aspirate, tilt the plate and aspirate from the well wall to avoid disrupting the gels.
8. Wash the gels by gently adding some **warm sterile PBS** to each well with a serological pipette.
    a. Remember to add the PBS by sliding it down the walls of the wells.
9. Gently aspirate the PBS.
10. Gently add fresh medium to each well with a serological pipette.

### Organoid Passaging (6-well plates)

#### Materials

- Cultrex RGF Basement Membrane Extract, Type 2, Pathclear (3533-005-02).
- Bucket with ice.
- Y-27632 10 mM ($F_c$ 5 µM).
- 1X Sterile PBS [-] $CaCl_2$ [-] $MgCl_2$.
- M1 and M2.
- 15-mL PP snap cap Falcon tubes (1 for the gel, 1 for the cold PBS, one for each sample).
- 50-mL Falcon tubes.
- 6-well culture plates Nunc Cell-Culture Treated (ThermoFisher: 140685).
- TrypLE Express.
- Cell scrapers.
- 10-mL serological pipettes.
- Electronic pipette and tips.
- Pipette set and tips.
- Sterile, Pasteur glass pipettes for the vacuum pump.

#### Important Notes

Organoids must not be over-pipetted. This is particularly important when organoids cultures are in early phases. Cells usually give rise to organoids in presence of other cells. In early phases of culture development, you might notice that the organoid growth is very slow and organoid are scarce. In such case, do not try to expand the number of wells cultured – instead, try to concentrate the organoids in a smaller amount of gel and make the culture denser. HGSC organoids show preference for growth in high-density culture. Once the culture is dense, you will notice that passaging/expansion ratio and time between passages become stable.

#### Instructions

##### *Early preparation*

1. Take the plates out from the incubator, observe them under the microscope to make sure that the organoids reached the desired size/density for passaging and no contamination is present.
2. Place the new 6-well plates in the incubator for at least 30 minutes.
    a. The BME should polymerize quickly in a warm plate, so that most cells don't attach to the plastic.
3. Aliquot some PBS in a 50 mL Falcon tube.
4. Place 2 snap cap tubes in the ice bucket (1 for the cold PBS and 1 for the gel).
5. Aliquot some PBS in 1 of the snap cap tubes and place it back on ice.

##### *Cell harvest*

6. Aspirate the media from all the wells.
7. Wash the gels with room-temerature sterile PBS with a serologic pipette.
8. Add 2 mL of TrypLE Express per well.
9. Scrape the gels off and detach them from the plate with a cell scraper.
    a. Remember to use a different cell scraper for each sample.
10. With the P1000, vigorously pipette the gels **(4-7 times depending on the cell density)** until disrupted while rinsing the whole well. The gels must be disrupted so that they digest well in TrypLE Express.
    a. In case of a low cell density, pre-wet the pipette tip with TrypLE Express (by aspirating and dispensing back to the bottle)
11. **Incubate the plate for 15' in the 37 degrees incubator**. The organoids must not be incubated with TrypLE Express longer than 25-30 minutes because organoids could be over-digested as well.

##### *During the 15-minute break prepare the gel*

12. Prepare the gel as described in previous sections.

*After the 15-minute break*

13. Take the plates out of the incubator (one at a time).
14. Take the P1000 and pre-wet the pipette tip with TrypLE Express.
15. Transfer the cells to a 15-mL snap cap Falcon tube. Try to get most of the gel the first time. Rinse the well with the leftover liquid the second time.
16. Add 1 mL of PBS to each well to rinse it nad transfer everything to the same tube in order to harvest the maximal number of organoids.
17. Spin down the organoids (300G, 5').
18. Gently aspirate the supernatant.
    a. Start by aspirating the bubbles on the surface of the supernatant, as the might disrupt the pellet.
19. Gently mix the BME solution with the P1000.
20. Add desired amount of the gel solution (200 µL + 50 µL extra volume per 10 droplets in a single well of 6-well plate. The more wells you seed, the less extra volume per well you will need) to the cell pellet and re-suspend it until obtaining a homogenous solution. <u>For delicate samples, pipette 10-12 times and around 14-18 for the sturdy ones.</u>
    a. The thicker the gel, the less accurate the volume is.
21. Take a new 6-well plate out of the incubator.
22. Seed the gels with the electronic pipette <u>(20 µL/droplet x 10 droplets/well).</u>
    a. Be careful not to place the gels neither too close to each other nor to the walls.
    b. Try to use as much of the organoid suspension as possible (e.g. seed the dead volume in the stepper pipette as well) – this is especially important when organoids are scarce in early culture development phase.

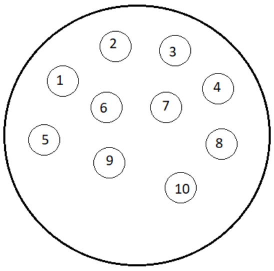

23. Describe the plates: patient ID, sample, passage, medium, and date.
24. <u>Incubate the plates for 45' in the incubator</u>.

*During the 45'-minute break*

1. Aliquot the desired amount (3 mL/well) of M1 and M2 media in Falcon tubes.
2. Add Y-27632 ($F_c$ 5µM) and mix.
3. Warm up the media at approx. 37ºC by placing the Falcon tubes in the water bath for approx. 10-15'.

*After the 45'-minute break*

25. <u>Carefully</u> add medium to the plates with a serologic pipette (do not pipette the medium directly on the gel domes – instead, dispense against the well wall, as the gels are delicate and disrupted easily).
26. Assess the seeding density and cell morphology under the microscope (it is important to observe every culture in order to adjust the passaging ratio for each one separately)
27. Place the plates in the incubator.

### Organoid Cryopreservation and Biobanking

#### Materials

- Pipette set and tips.
- Cryovials for cell storage.
- Printed labels.
- Stem-Cellbanker (#11890, Amsbio (DMSO-free freezing solution. When using it, samples can be transferred to -80 right away. Samples can be transferred to the nitrogen storage tank 24 h after being frozen).

#### Instructions

1. Label to the cryovials.

2. After harvesting the cells and aspirating the supernatant (as decribed in Organoid Passaging), add Stem-Cellbanker to the pellets as following:
    a. If the pellet is from 1 well:
        i. Add 1 mL of Stem-Cellbanker, take up the pellet, and transfer it to a cryovial.
        ii. Transfer the rest of the Stem-Cellbanker to the cryovial.
        iii. Should there be more cells in the tube, rinse it with 250 µL of Stem-Cellbanker.
        iv. Re-suspend the cells in the cryovial using a P1000 pipette (5-7 times) to reach a homogenous solution without large pellet fragments (these do not freeze well). Do not over-pipette the organoids.
    b. If the pellet is from more than 1 well:
        i. Add 1 mL (per each prepared cryovial) of Stem-Cellbanker to the cell pellet.
        ii. Re-suspend to achieve a homogenous cell suspension, and aliquot it in the cryovials.
3. Place the samples at the -80 freezer.

IMPORTANT: When thawing a cryopreserved organoids, reduce the initial passaging ratio by ½ (e.g. for organoid culture passaged at 1:4 ratio, seed the cryopreserved material from a single well to 2, instead of 4 wells). You will notice that there is an increased amount of dead cells after thawing and the organoid growth might be initially slower. This is normal and the culture should stabilize after 1-2 passages, returning to the old growth/passaging ratio.

### Media Preparation

#### Materials

- Advanced DMEM/F12 (1X) (+NEAA; +sodium pyruvate; -L-Glutamine).
- Supplements, growth factors, and small molecule inhibitors.
- 1 15-mL Falcon tube.
- 1 50-mL Falcon tube.
- Plastic spoon and spatula.
- 250-mL sterile Corning bottle.
- Pipette set and tips.

#### Important Notes

Small molecules in DMSO can be refrozen up to 3 times.
FGF-4, FGF-10, EGF, neuregulin-1 and hormones cannot be re-frozen.

#### Instructions

##### Medium 1

1. Thaw the reagents in advance.
2. Weight the N-Acetyl-L-Cysteine and the Nicotinamide powders.
3. Spin down all the aliquots beforehand.
4. Add the different reagents into a bottle of Advanced DMEM/F12 Medium to prepare M1.
    a. HEPES, GlutaMAX, and B-27 can be poured directly into the medium flask. Tubes must be rinsed with the medium
    b. Add B-27 before the growth factors, as it contains BSA, which prevents growth factor molecules from attaching to plastic.
    c. N-Acetyl-L-Cysteine and Nicotinamide can be dissolved by adding some media into the Falcon tubes and mixing.
        i. N-Acetyl-L-Cysteine takes longer to dissolve, so repeat the operation as many times until there aren’t any crystals left in the Falcon tube.
5. Once M1 is prepared, mix it thoroughly.

##### Medium 2

6. Pour the desired amount of M1 into a sterile plastic bottle.
7. Add EGF, Neuregulin-1, Forskolin, and hydrocortisone.
8. Mix thoroughly and store it in the fridge.

| Medium 1 | Stock concentration | | Final concentration | | Initial Volume | |
|---|---|---|---|---|---|---|
| | Value | Units | Value | Units | Amount | Units |
| Advanced DMEM/F12 (1X) (+NEAA; +sodium pyruvate; -L-Glutamine)F | - | - | - | - | 500 | mL |
| PrimocinFz | 50 | mg/mL | 100 | µg/mL | 1 | mL |
| HEPESFz | 1 | M | 10 | mM | 5 | mL |
| GlutaMAXFz | 100 | X | 1 | X | 5 | mL |
| N-Acetyl-L-CysteineF | 163,1951 | g/mol | 1 | mM | 0,08159755 | g (+4mg) |
| NicotinamideRT | 122,12 | g/mol | 5 | mM | 0,3053 | g (+4mg) |
| B-27Fz | 50 | X | 1 | X | 10 | mL |
| β-estradiolFz | 10 | mM | 100 | nM | 5 | µL |
| SB202190Fz | 10 | mM | 0,5 | µM | 25 | µL |
| A83-01Fz | 10 | mM | 0,5 | µM | 25 | µL |
| FGF-4Fz | 100 | µg/mL | 10 | ng/mL | 50 | µL |
| FGF-10Fz | 100 | µg/mL | 10 | ng/mL | 50 | µL |
| ***Y-27632 | 10 | mM | 5 | µM | 250 | µL |
| **Medium 2** | **Stock concentration** | | **Final concentration** | | **Initial Volume** | |
| | Value | Units | Value | Units | Amount | Units |
| Medium 1 | - | - | - | - | 300 | mL |
| Neuregulin-1Fz | 50 | µM | 5 | nM | 30 | µL |
| EGFFz | 100 | µg/mL | 5 | ng/mL | 15 | µL |
| ForskolinFz | 10 | mM | 5 | µM | 150 | µL |
| HydrocortisoneFz | 2 | mg/mL | 500 | ng/mL | 75 | µL |

**Key**

| | |
|---|---|
| Blue: prepare 5-mL aliquots in advance | RT room temperature |
| Green: powder | F fridge |
| Red: mark and refreeze after use. | Fz -20 ºC |
